# Dynamic Auditory Remapping Across the Sleep-Wake Cycle

**DOI:** 10.1101/2021.02.16.431383

**Authors:** Anat Arzi, Caterina Trentin, Annamaria Laudini, Alexandra Krugliak, Dritan Nikolla, Tristan Bekinschtein

## Abstract

In a single day we transition from vigilant wakefulness to unconscious sleep and dreaming, undergoing diverse behavioural, physiological and neural changes. While during the awake state, exogenous stimuli and endogenous changes lead to sensory reorganisation, this remapping has not been charted throughout the sleep-wake cycle. We recorded neural activity in response to a range of tones using electroencephalography during a full night’s sleep, and examined whether auditory responses become more similar, dissimilar or remain unchanged between wakefulness, non-rapid (NREM) and rapid eye movement (REM) sleep. We found that neural similarities between pairs of auditory evoked potentials differed by conscious state in both early and late auditory processing stages. Furthermore, tone-pairs neural similarities were modulated by conscious state as a function of tone frequency, where some tone-pairs changed similarity between states and others continued unaffected. These findings demonstrate a state-, stimulus- and time-dependent functional reorganization of auditory processing across the sleep-wake cycle.

## Introduction

From the first day of life (Fifer et al., 2010) and throughout adulthood (Clancy et al., 2019; Danielson et al., 2016; Fritz et al., 2003; Heed et al., 2015) the neural basis of sensory processing is dynamically shaped by the interaction between our internal state and the environment. Reorganization of sensory, motor and cognitive functions, also termed remapping (Ramachandran et al., 1992), is expressed in the nervous system in different forms and time scales. Somatosensory remapping, in which the cortical area corresponding to a specific body part becomes responsive to stimuli applied to different body areas, is evident following permanent neural damage such as amputation. (Buonomano and Merzenich, 1998; Merzenich et al., 1984; Pons et al., 1991; Ramachandran et al., 1992). In addition, recovery from stroke induces remapping of language functions between hemispheres and of motor functions beyond motor areas (Carmichael, 2003), retinal lesions promote remapping of the retinotopic organization of the neuronal receptive fields in the visual cortex (Dumoulin and Knapen, 2018), and partial deafness results in cortical reorganisation of frequency representations (King and Moore, 1991). Remapping also take place in healthy animals and humans, when applying prismatic spectacles temporarily shifting the visual map (Bultitude et al., 2013; Linkenhoker and Knudsen, 2002), by artificially adjoining fingers leading to reorganisation in the somatosensory cortex (Buonomano and Merzenich, 1998; Kolasinski et al., 2016), or following repetitive transcranial magnetic stimulation to the primary motor cortex inducing widespread changes in the motor system (Lee et al., 2003). Moreover, remapping naturally occurs without such manipulations. For example, place cells in the hippocampus that preferentially fire to distinct regions of a spatial environment remap their spatial preference in response to changes in shape, light, colour and familiarity of the environment (Alexander et al., 2016; Fyhn et al., 2007; Geva-Sagiv et al., 2016; Hayman et al., 2003; Jeffery, 2011; Leutgeb et al., 2005; Moser et al., 2014), and grid cells’ maps in the entorhinal cortex are dynamically restructured by cognitive factors or running speed (Boccara et al., 2019; Butler et al., 2019; Low et al., 2020; Quiroga, 2019) . Furthermore, not only changes in the environment but also endogenous signals such as vestibular cues, motivation, oxytocin or norepinephrine levels elicit functional reorganization, leading to differential neural responses to identical sensory stimuli (Doboli et al., 2003; Grella et al., 2019; Knierim et al., 1998; Marlin et al., 2015). The fact that transitioning between wakefulness and different sleep stages involves substantial external and internal changes (Fuller et al., 2006; Jones, 2005) suggests that sleep may lead to sensory remapping, however, it is unknown whether sensory maps remain stable across the sleep-wake cycle or whether sleep causes functional sensory reorganisation.

There is a wide consensus that some degree of sensory processing persists during sleep (Andrillon and Kouider, 2020; Arzi et al., 2012, 2014; Atienza et al., 2001; Canales-Johnson et al., 2020; Chennu and Bekinschtein, 2012; Hennevin et al., 2007; Velluti, 1997). Neuroimaging and electrophysiology studies in humans (Atienza et al., 2001; Bastuji and García-Larrea, 1999; Colrain and Campbell, 2007; Czisch et al., 2002, 2009; Portas et al., 2000; Schabus et al., 2012; Wilf et al., 2016) and animals (Edeline et al., 2001; Issa and Wang, 2008; Nir et al., 2015; Sela et al., 2020) demonstrate clear brain activity in response to sensory stimuli in sleep. However, it remains unclear precisely how sensory-related neural activity changes during sleep. Diverse results are reported for the degree of modulation of sensory responses between wakefulness and sleep, with findings of enhanced (Colrain and Campbell, 2007; Hall and Borbely, 1970; Nicholas et al., 2006; Yang and Wu, 2007), reduced (Brugge and Merzenich, 1973; Czisch et al., 2002, 2004; Edeline et al., 2001; Murata and Kameda, 1963) or preserved responses during sleep (Edeline et al., 2001; Issa and Wang, 2008; Nir et al., 2015; Peña et al., 1999). Furthermore, several studies suggest that even within the same experiment, the degree of response modulation between conscious states is not consistent across stimuli and may depend on stimulus intensity or type (Castro-alamancos, 2004; Issa and Wang, 2011; Lustenberger et al., 2018; Portas et al., 2000; Sharon and Nir, 2018; Tlumak et al., 2012). This suggests that the magnitude and direction of changes in sensory responses may vary depending not only on the sleep state, but also on the properties of the stimuli. Yet, the investigation of sensory processing during sleep has focused on the modulation of neural responses to either a specific stimulus or a set of stimuli, while neglecting the investigation of the modulation of the neural similarity between responses to different stimuli. In other words, how the degree of similarity between sensory responses is altered between wakefulness and sleep and between sleep stages: do brain responses to sensory stimuli become more similar to one another, more different from each other or do they remain unchanged across the sleep-wake cycle?

To address this question, we employ the auditory frequency axis as the preferred mode of input for unconscious states (Chennu and Bekinschtein, 2012). Audition is particularly attractive to study sensory reorganisation during sleep for several reasons: first, tones can be easily presented during sleep without waking the participant; second, tones presentation can be done with high temporal precision and short inter-stimulus intervals, enabling the collection of thousands of tone repetitions per participant; third, the auditory frequency axis offers a clear measure of distance between the physical properties of pure tones; fourth and last, the neural representations of pure tones during the awake state are well-described (Saenz and Langers, 2014; Schnupp et al., 2011; Su et al., 2014). Thus, the auditory frequency axis is an ideal experimental model system to study functional sensory reorganisation across the sleep-wake cycle. Here, we recorded brain activity in response to a series of pure tones in the middle part of the human audible frequency range (Fig. 1), during wakefulness and throughout a full night’s sleep using high-density electroencephalography (EEG). Given the vast internal changes between wakefulness and different sleep stages, we hypothesised that the organisation of the auditory frequency map is modulated by conscious state in a sleep-stage dependent manner. Furthermore, we also hypothesised that the shape of the reorganisation depends on tone frequency. To test these hypotheses, we assessed how similarities between neural responses to tone-pairs change across the sleep-wake cycle. Specifically, we measured neural similarities between tone-pairs in wakefulness, light (N2) and deep (N3) non-rapid eye movement (NREM) sleep, and rapid eye movement (REM) sleep, and found that they dynamically change in time between and also within conscious states. Moreover, tone-pairs neural similarities were modulated in an uneven manner across conscious states as a function of tone frequency. These findings are consistent with our hypothesis of sleep-induced remapping, indicating that the functional sensory organisation is state-, stimulus- and time-dependent.

**Fig. 1:**
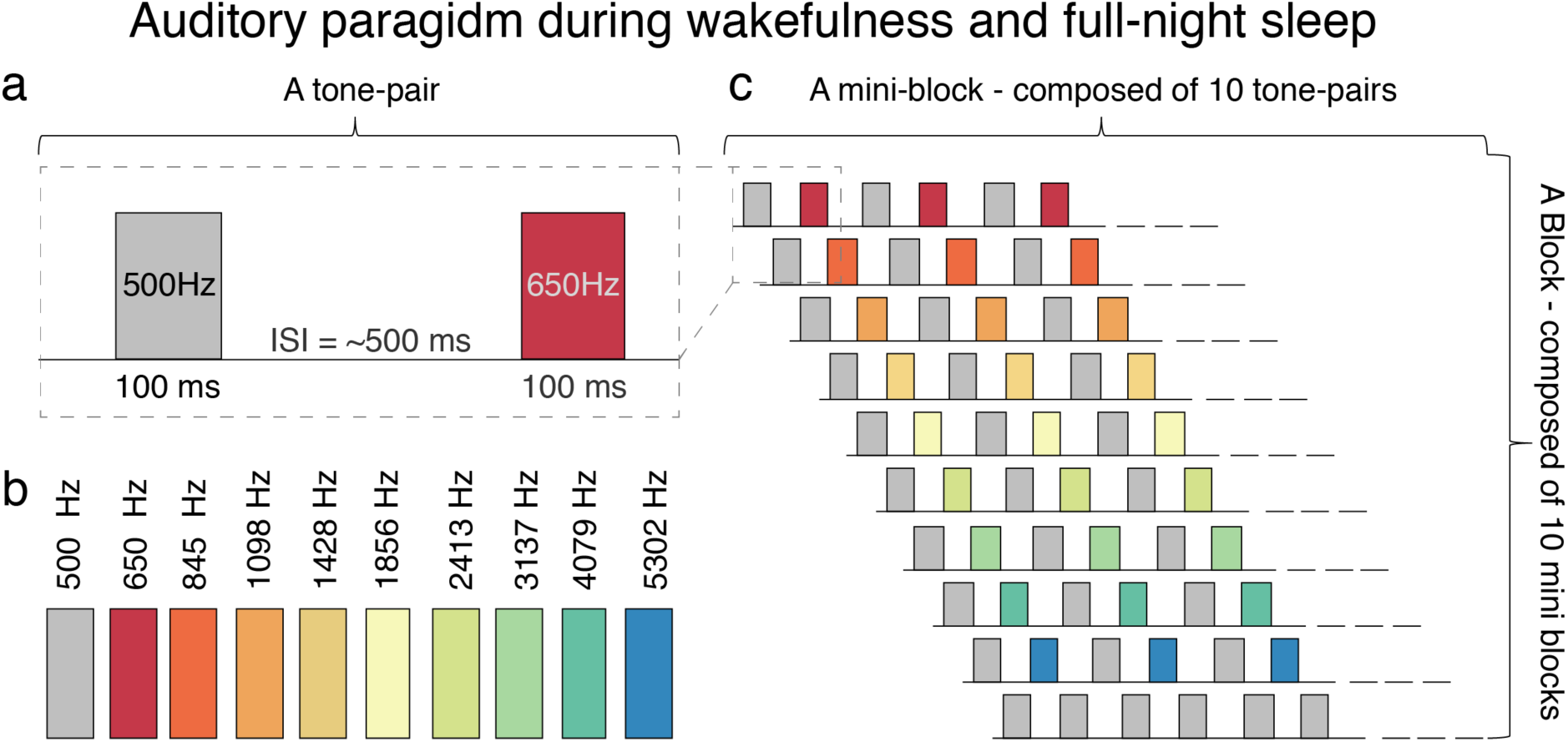
Experimental design. A diagram of the auditory experimental paradigm based on tone-pairs. **a)** In each tone-pair, the first tone was an ‘Adaptor’ tone (500 Hz, grey) which was presented to create a common context for all tones, and was followed by one of nine pure tones or by the adaptor **b)** The tones presented were spaced by 30% from one another: 500 (adaptor tone), 650, 845, 1098, 1428, 1856, 2413, 3137, 4079 and 5302 Hz. **c)** In a mini-block, each tone-pair was repeated for 10 times and 10 mini-blocks presented in a random order created a *block*. A wakefulness session was composed of 24 blocks and lasted approximately an hour. In a sleep session, the number of blocks depended on sleep duration (Tables 1 and 2).

**Table 1:** Number of trials averaged across participants.

|  | **T650** | **T845** | **T1098** | **T1428** | **T1856** | **T2413** | **T3137** | **T4079** | **T5302** | **Total** |
| --- | --- | --- | --- | --- | --- | --- | --- | --- | --- | --- |
| **Awake** | 217.69 ± 9.88 | 217.56 ± 10.32 | 217.39 ± 9.43 | 217.89 ± 10.30 | 217.64 ± 9.96 | 217.58 ± 9.37 | 217.44 ± 9.14 | 217.36 ± 9.71 | 217.72 ± 9.59 | 1958.28 ± 86.73 |
| **N2 sleep** | 627.72 ± 200.78 | 633.44 ± 200.57 | 626.83 ± 201.80 | 624.64 ± 195.21 | 621.81 ± 194.78 | 614.0 ± 198.48 | 602.81 ± 191.23 | 616.44 ± 203.21 | 627.06 ± 206.97 | 5594.75 ± 1776.20 |
| **N3 sleep** | 167.31 ± 78.03 | 161.94 ± 73.60 | 160.64 ± 74.08 | 155.03 ± 67.44 | 158.25 ± 69.20 | 155.75 ± 69.22 | 151.03 ± 70.67 | 157.5 ± 73.47 | 158.72 ± 74.20 | 1426.17 ± 638.79 |
| **REM sleep** | 225.5 ± 115.68 | 226.19 ± 113.69 | 228.94 ± 116.11 | 221.17 ± 106.59 | 224.58 ± 110.17 | 229.33 ± 119.29 | 226.06 ± 113.86 | 220.75 ± 112.61 | 223.08 ± 112.62 | 2025.61 ± 1013.86 |

Averaged ± SD number of trials across participants for each tone in wakefulness and sleep.

**Table 2:** Sleep architecture.

| Awake | N1 sleep | N2 sleep | N3 sleep | REM sleep | Total sleep time | Time in bed |
| --- | --- | --- | --- | --- | --- | --- |
| 137.46 ±<br>52.49 | 26.42 ±<br>19.41 | 161.35 ±<br>48.76 | 56.42 ±<br>25.36 | 51.43 ±<br>25.26 | 295.61 ±<br>68.87 | 433.07 ±<br>61.92 |

Averaged ± SD time (in minutes) spent in each sleep stage

## Results

### Auditory evoked responses are state-, time- and stimulus-dependent

To test the hypothesis that transitions between wakefulness and different sleep stages (N2, N3 and REM sleep) induce functional sensory reorganisation, we recorded brain activity using high-density EEG during wakefulness and full-night sleep in response to nine pure tones (650, 845, 1098, 1428, 1856, 2413, 3137, 4079 and 5302 Hz; Fig. 1), acquiring thousands of trials per conscious state (Table 2).

First, we examined the auditory evoked related responses (ERP) dynamics. ERPs averaged across tones were compared between wakefulness (W), N2, N3 and REM sleep using cluster permutation analysis. ERPs waveform showed greater voltage negativity (N100) in wakefulness and greater voltage positivity (P200) during sleep (Fig. 2), with differences observed in three main time windows (**W-N2**: 20-108 ms, p = 0.0013, effect size Wr = 0. 859; 124-316 ms, p = 0.0001, effect size Wr = 0.819; 348-448 ms, p = 0.152, effect size Wr = 0.395; Horizontal blue-yellow line; **W-REM**: 16-96 ms, p = 0.0003, effect size Wr = 0.872; 136-344 ms, p = 0.0001, effect size Wr = 0.820; Horizontal blue-green line; **W-N3**: 28-100 ms, p = 0.0039, effect size Wr = 0.848; 128-316 ms, p = 0.0002, effect size Wr = 0.807; 364-448 ms, p

**Fig. 2:**
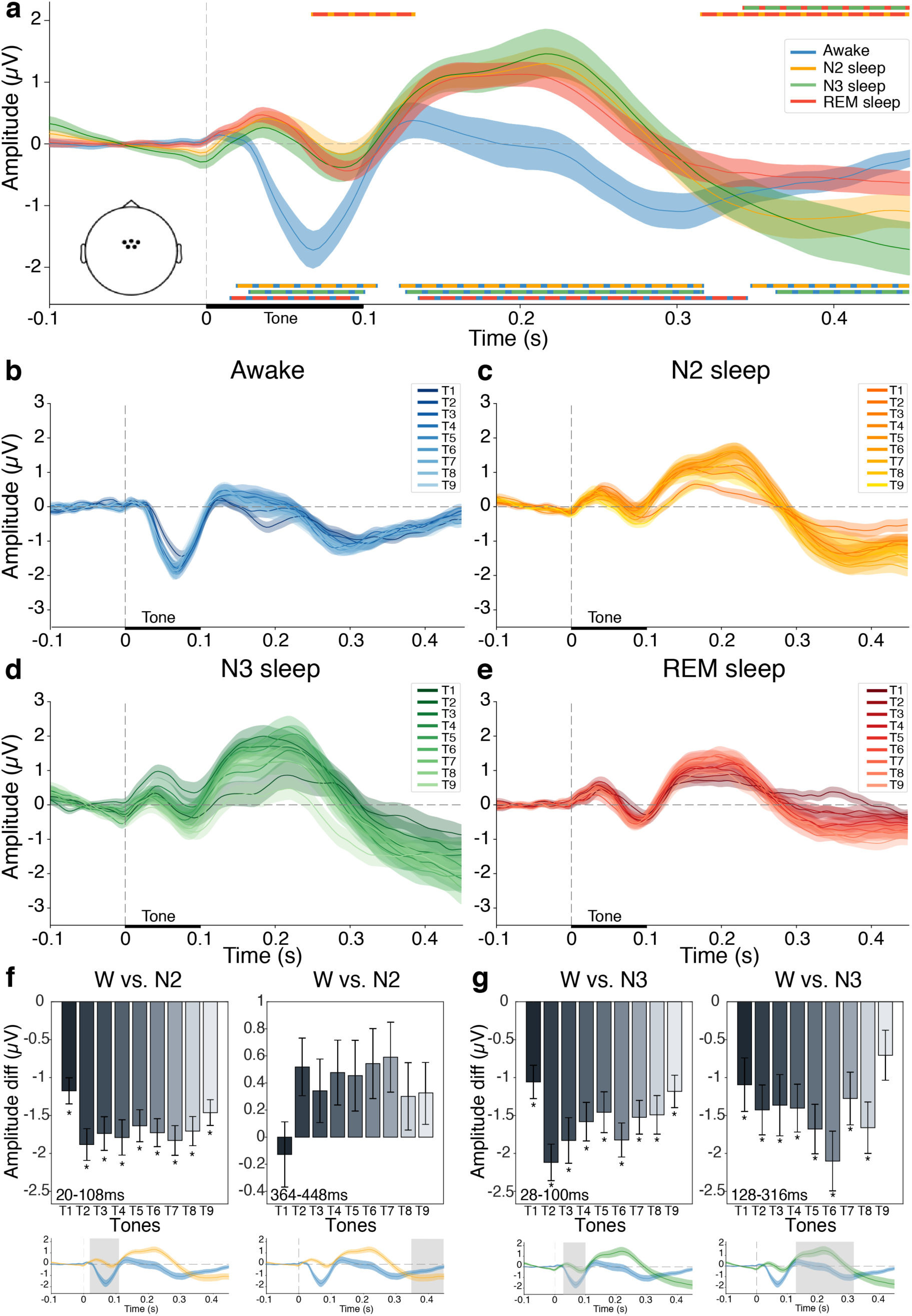
Auditory evoked responses are state and stimulus dependent. a) Auditory evoked potentials averaged across participants and tones during wakefulness (blue), N2 sleep (yellow), N3 sleep (green) and REM sleep (red). The dashed vertical line denotes tone onset (duration 100 ms, black horizontal line). The horizontal color lines denote significant differences between pairs of states showing greater voltage negativity (N100) in wakefulness and greater voltage positivity (P200) during sleep (Bottom lines: Awake vs. N2, blue-yellow line; Awake vs. N3, blue-yellow line; Awake vs. REM, blue-green line) as well as differences between NREM and REM sleep (Top lines: N2 vs. REM, yellow-red line; N3 vs REM, green-red line;) **b-e)** Auditory evoked potentials for each tone frequency during **b)** wakefulness **c)** N2 sleep **d)** N3 sleep **e)** REM sleep **f-i)** Interactions between conscious states and tone frequency were found between **f)** wakefulness and N2 at 20-108ms (p = 0.02, left) and at 28-100ms (p = 0.024, right) and between **g)** wakefulness and N3 at 28-100ms (p = 0.01, left) and at 128-316ms (p = 0.036, right). Bars denotes the mean and error bars the S.E.M of 12ms at the centre of each cluster. Each cluster time and duration is indicated by a shaded area on the ERP plot below the relevant bar plot. * denote a significant difference in amplitude between pairs of states per tone, FDR-corrected for multiple comparisons. T1 =650Hz, T2 =845Hz, T3 =1098Hz, T4 =1428Hz, T5 =1856Hz, T6 =2413Hz, T7 =3137Hz, T8 =4079Hz and T9 =5302Hz.

= 0.0157, effect size Wr = 0.456; Horizontal blue-red line; Fig. 2 and Table 1). Smaller and delayed ERPs differences were detected between sleep stages, specifically between NREM and REM sleep (**N2-REM**: 68-132 ms, p = 0.043, effect size Wr = 0.547; 316-448, p = 0.0012, effect size Wr = 0.393; Horizontal yellow-red line; **N3-REM**: 348-448 ms, p = 0.009, effect size Wr = 0.427; Horizontal green-red line; Fig. 2). No detectable changes in ERPs were found between NREM sleep stages (**N2-N3**: all p’s > 0.13; Fig. 2). To examine whether the observed differences between conscious states were influenced by tone frequency, we applied a linear mixed-effects modelling analysis including all tones and pairs of conscious states in each one of the 11 identified clusters. Interactions between *conscious state* and *tone* were found between wakefulness and NREM sleep, in both N2 sleep (**W-N2**: 20-108 ms, p = 0.02, Fig. 2f; 364-448 ms, p = 0.024, Fig. 2i) and N3 sleep (**W-N3**: 28-100 ms, p = 0.01, Fig. 2g; 128-319 ms p = 0.036; Fig. 2h). These interactions reflect smaller ERP differences between wakefulness and NREM sleep stages in low and high tone frequencies and larger ones in mid-frequencies. These findings indicate an uneven reshaping of the auditory response by conscious state and support the hypothesis of stimulus-dependent modulation of auditory processing across the sleep-wake cycle.

### The magnitude of the auditory neural similarities are state-, time- and stimulus-dependent

To further the understanding how conscious state modulates auditory processing, we examined the relationship between neural responses to pairs of pure tones in wakefulness, NREM and REM sleep. We quantified the neural similarity between ERP-pairs using Spearman’s correlation, and termed this distance measure *Tone Similarity Index* (see methods). Measuring the Tone Similarity Index along the auditory processing time course (i.e., the 450ms following stimulus presentation) averaged across all tone-pairs combinations, we discovered that it is dynamically changing over time (Fig. 3a). Using cluster permutation analysis, we identified two main time windows in which the Tone Similarity Index exhibits a distinctive pattern: An early (tw1 = 12-236 ms), and a late (tw2 = 256-448 ms) processing time windows, likely representing different auditory processing stages. Next, to characterise how the neural similarity between tone-pairs is changing in time between conscious states, we applied linear mixed-effects modelling. Specifically, linear mixed-effects modelling was used to understand how the Tones Similarity Index in each participant was influenced by conscious state (‘Wakefulness’, ‘N2’, ‘N3’, and ‘REM’), auditory processing time window (‘Early’ and ‘Late’), and tone-pair (36 pairwise combinations, created from nine tone frequencies). The model that best fitted the data was one with *participants* as random effect, and *conscious state*, *time window* and *tone-pair* as fixed effects (Table 3). Results showed a main effect of *state* (F(3,10150) = 53.26, p < 0.0001, *η*^2^ = 0.02; Fig. 3a), indicating that conscious state has an effect on auditory processing, a main effect of *tone-pair* (F(35, 10150) = 42.35, p < 0.0001, *η*^2^ = 0.13), reflecting tone-pair specific similarity magnitudes (Fig. 2b-e, Fig 3b-c), and a main effect of *time window* (F(1, 10150) = 1186.03, p < 0.0001, *η*^2^ = 0.10; Fig. 3 a-c), in line with the observed dynamic changes in time of the Tone Similarity Index. In addition, interactions between *state* and *time window* (F(3, 10150) = 810.1, p < 0.0001, *η*^2^ = 0.19), *state* and *tone-pair* (F(105, 10150) = 2.79, p < 0.0001, *η*^2^ = 0.03), and *time window* and *tone-pair* (F(35, 10150) = 2.31, p < 0.0001, *η*^2^ = 0.008) were observed. The reliable interactions indicate that the relationship between neural responses to pure tones changes differently between conscious state depending on the time window and tones frequency, with some tone-pairs showing a significant change in similarity between states while others remaining unchanged (Fig. 4).

**Fig. 3:**
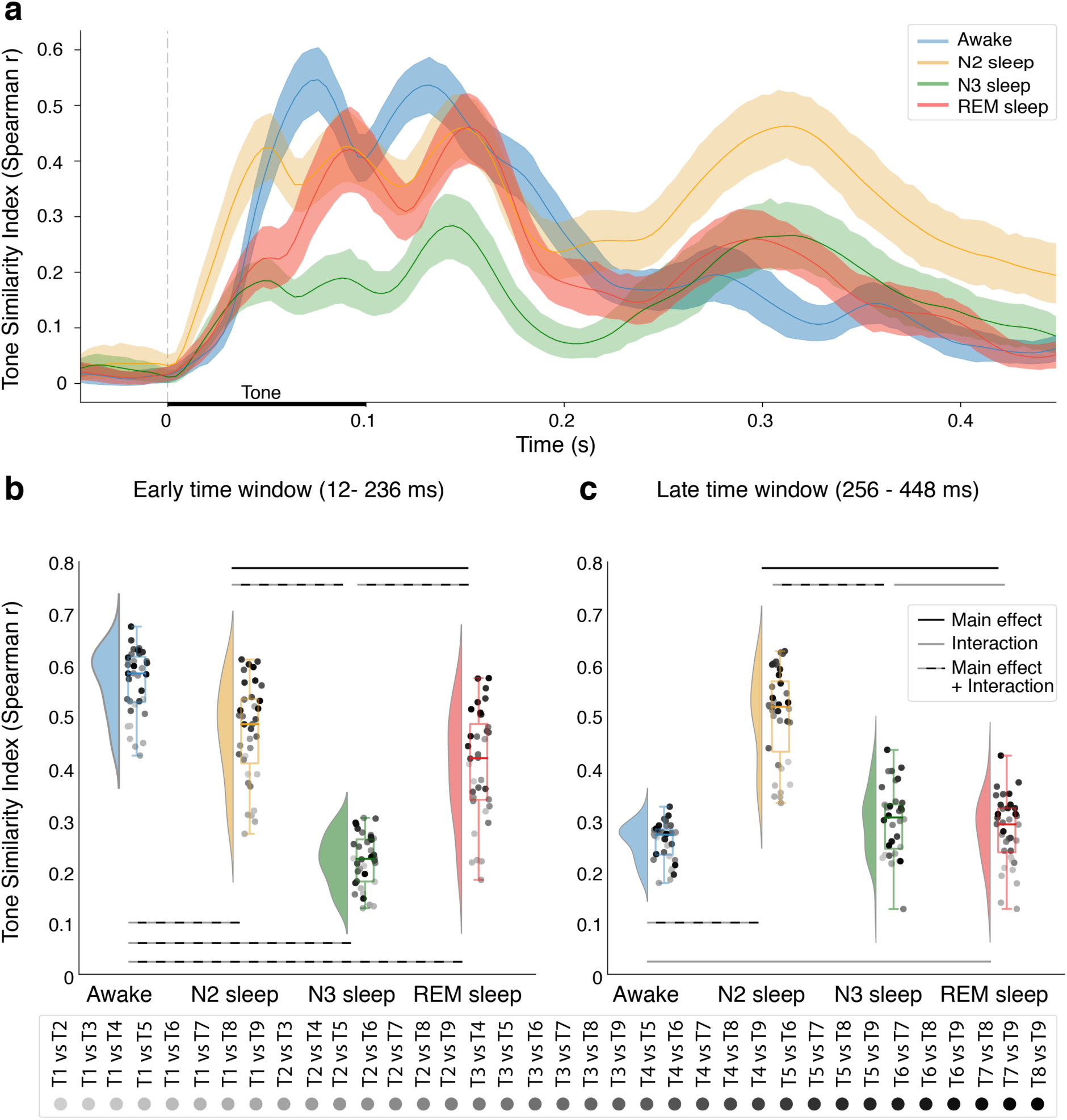
Similarity magnitude between auditory responses is state-, time- and stimulus-dependent. a) Auditory evoked responses similarity along the auditory processing time during wakefulness (blue), N2 sleep (orange), N3 sleep (green) and REM sleep (red) demonstrating a dynamic change in tone-pairs similarities across auditory processing time. **b-c)** Tone-pairs similarity in the **b)** early and **c)** late time windows showing reorganisation of tone-pairs similarities between states. Each dot represents a tone-pair; flat violin plots show the distribution; boxplot mid-line denotes the median; and the rectangle denotes the interquartile range (25th to the 75th percentiles). Horizontal black lines denote a main effect of conscious state; grey lines denote an interaction between conscious state and tone-pair (with no main effect of conscious state); black-grey lines denote a main effect of state and an interaction between conscious state and tone-pair.

**Fig. 4:**
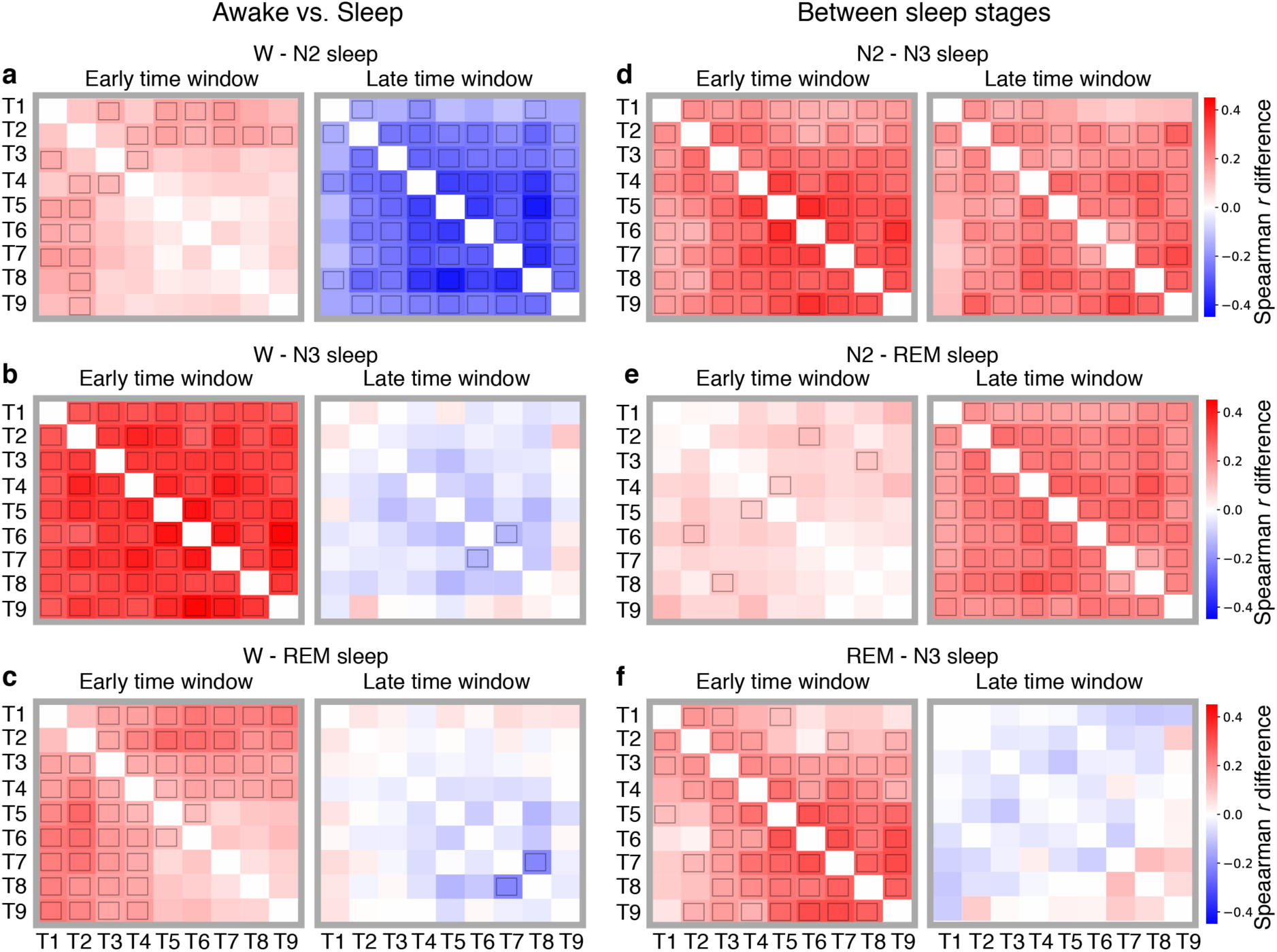
Auditory similarity magnitude differences between conscious states a-c) Differences in Representational Similarity Matrices (RSMs) between wakefulness and different sleep stages. Each matrix corresponds to the subtraction between pairs of similarity matrices (Fig. 5). **a)** RSMs difference between wakefulness and N2 sleep in the early (left) and late (right) time window. **b)** RSMs difference between wakefulness and N3 sleep in the early (left) and late (right) time window. **c)** RSMs difference between wakefulness and REM sleep in the early (left) and late (right) time window. **d-f)** Differences in Representational Similarity Matrices (RSMs) between wakefulness and different sleep stages. **d)** RSMs difference between N2 and N3 sleep in the early (left) and late (right) time window. **e)** RSMs difference between N2 and REM sleep in the early (left) and late (right) time window. **f)** RSMs difference between REM and N3 sleep in the early (left) and late (right) time window. Small black rectangle within each cell represent significant difference between states for a specific tone-pair, as obtained from planned comparisons following the computation of the linear mixed-effect models of tables 4 and 5 (see Methods).

**Table 3:**
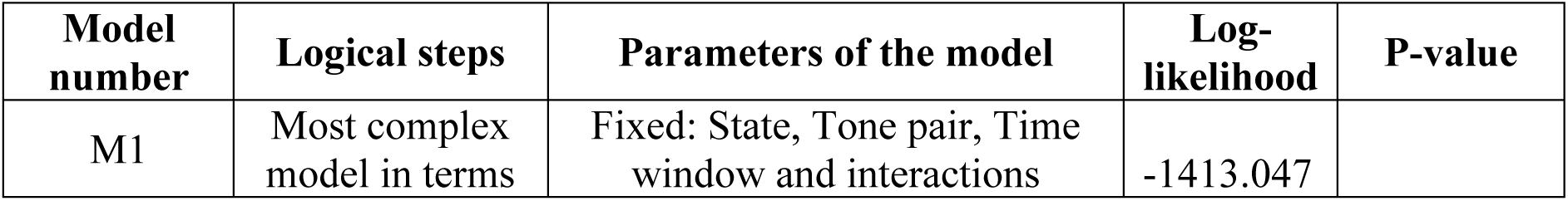
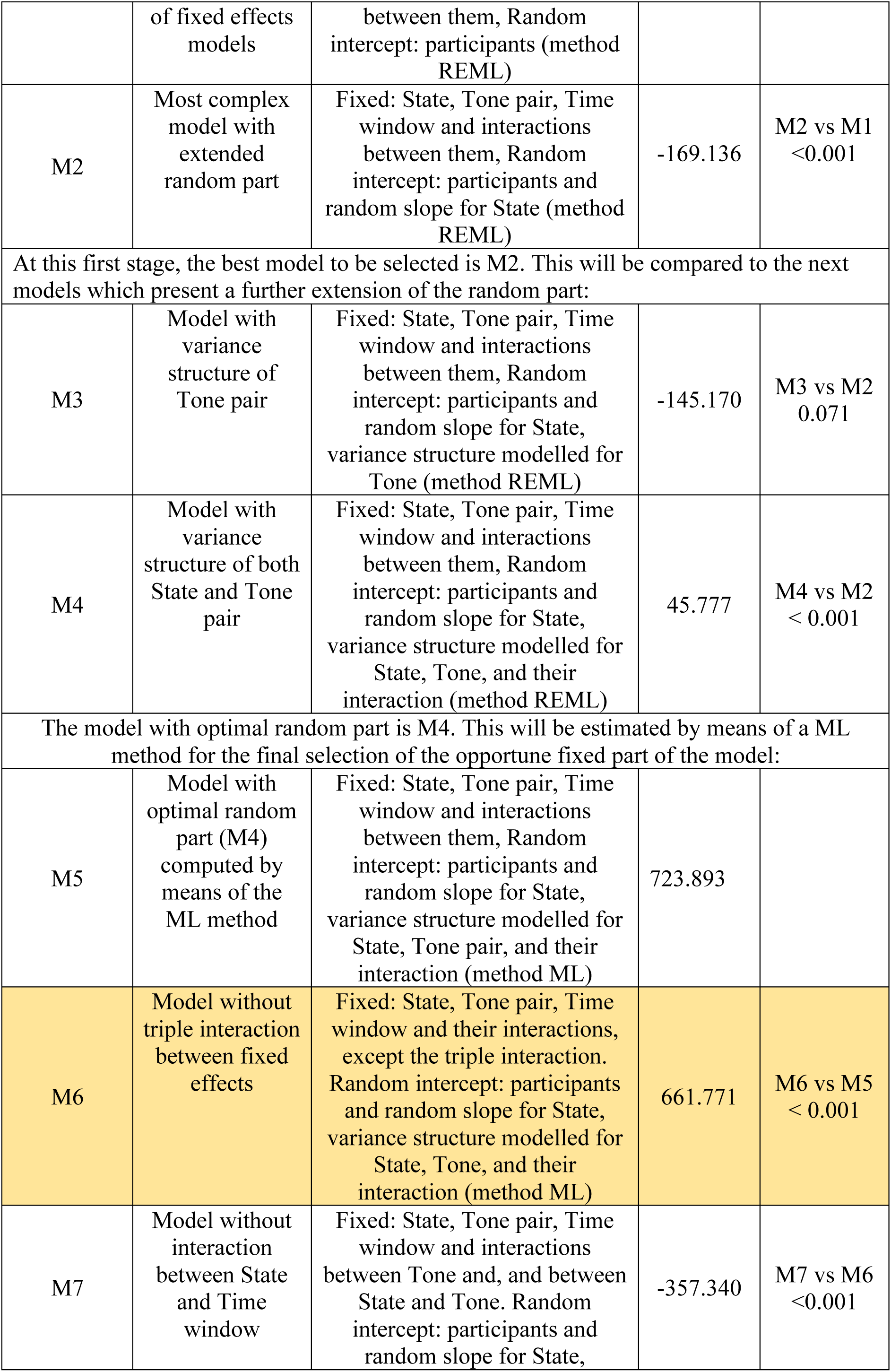
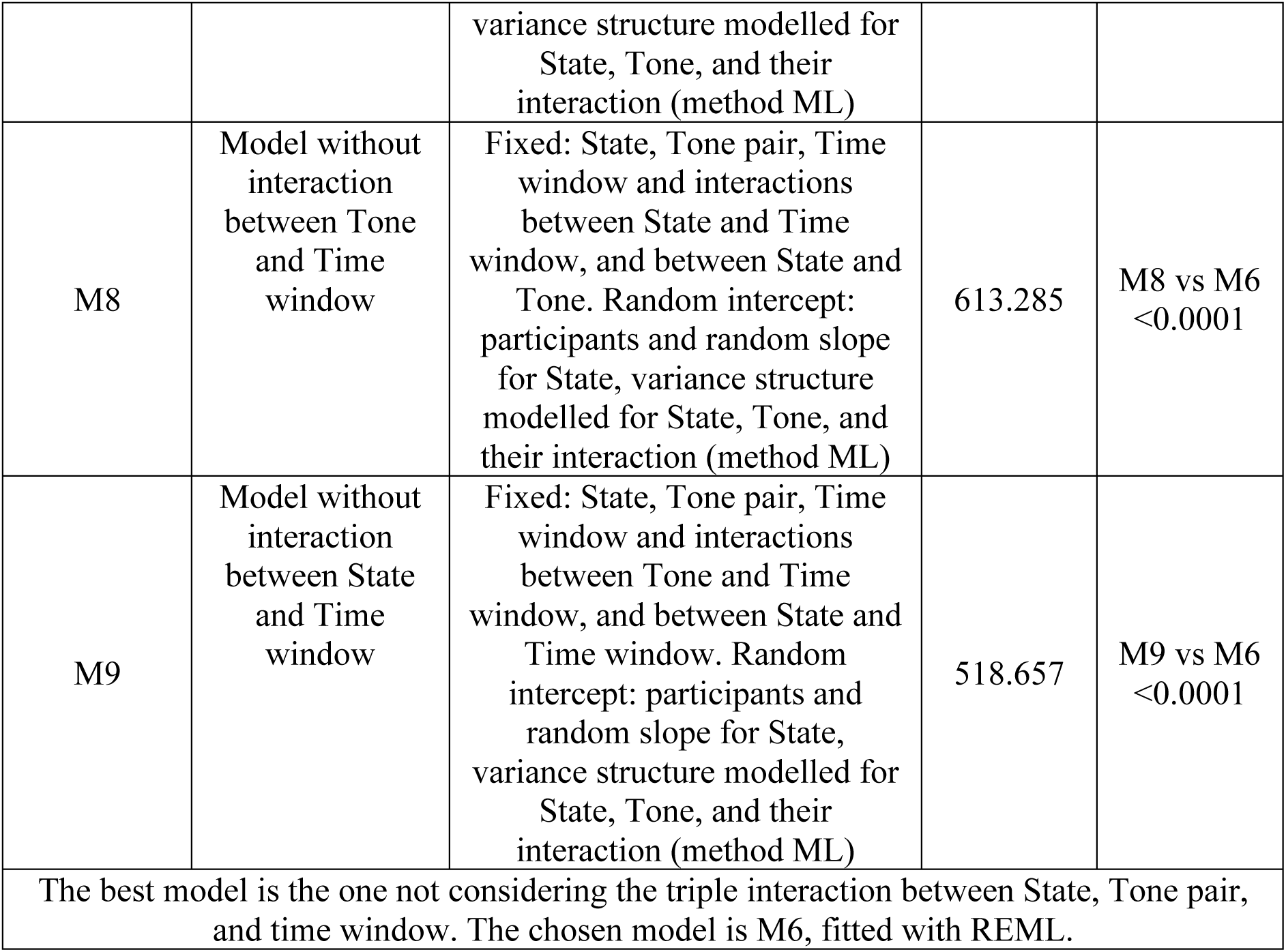
Omnibus linear mixed-effects models.

| Model number | Logical steps | Parameters of the model | Log-likelihood | P-value |
| --- | --- | --- | --- | --- |
| M1 | Most complex model in terms | Fixed: State, Tone pair, Time window and interactions | -1413.047 |  |
|  | of fixed effects models | between them, Random intercept: participants (method REML) |  |  |
| M2 | Most complex model with extended random part | Fixed: State, Tone pair, Time window and interactions between them, Random intercept: participants and random slope for State (method REML) | -169.136 | M2 vs M1 <0.001 |
| At this first stage, the best model to be selected is M2. This will be compared to the next models which present a further extension of the random part: |  |  |  |  |
| M3 | Model with variance structure of Tone pair | Fixed: State, Tone pair, Time window and interactions between them, Random intercept: participants and random slope for State, variance structure modelled for Tone (method REML) | -145.170 | M3 vs M2 0.071 |
| M4 | Model with variance structure of both State and Tone pair | Fixed: State, Tone pair, Time window and interactions between them, Random intercept: participants and random slope for State, variance structure modelled for State, Tone, and their interaction (method REML) | 45.777 | M4 vs M2 < 0.001 |
| The model with optimal random part is M4. This will be estimated by means of a ML method for the final selection of the opportune fixed part of the model: |  |  |  |  |
| M5 | Model with optimal random part (M4) computed by means of the ML method | Fixed: State, Tone pair, Time window and interactions between them, Random intercept: participants and random slope for State, variance structure modelled for State, Tone pair, and their interaction (method ML) | 723.893 |  |
| M6 | Model without triple interaction between fixed effects | Fixed: State, Tone pair, Time window and their interactions, except the triple interaction. Random intercept: participants and random slope for State, variance structure modelled for State, Tone, and their interaction (method ML) | 661.771 | M6 vs M5 < 0.001 |
| M7 | Model without interaction between State and Time window | Fixed: State, Tone pair, Time window and interactions between Tone and, and between State and Tone. Random intercept: participants and random slope for State, variance structure modelled for State, Tone, and their interaction (method ML) | -357.340 | M7 vs M6 <0.001 |
| M8 | Model without interaction between Tone and Time window | Fixed: State, Tone pair, Time window and interactions between State and Time window, and between State and Tone. Random intercept: participants and random slope for State, variance structure modelled for State, Tone, and their interaction (method ML) | 613.285 | M8 vs M6<br><0.0001 |
| M9 | Model without interaction between State and Time window | Fixed: State, Tone pair, Time window and interactions between Tone and Time window, and between State and Time window. Random intercept: participants and random slope for State, variance structure modelled for State, Tone, and their interaction (method ML) | 518.657 | M9 vs M6<br><0.0001 |
| The best model is the one not considering the triple interaction between State, Tone pair, and time window. The chosen model is M6, fitted with REML. |  |  |  |  |

**Table 4:**
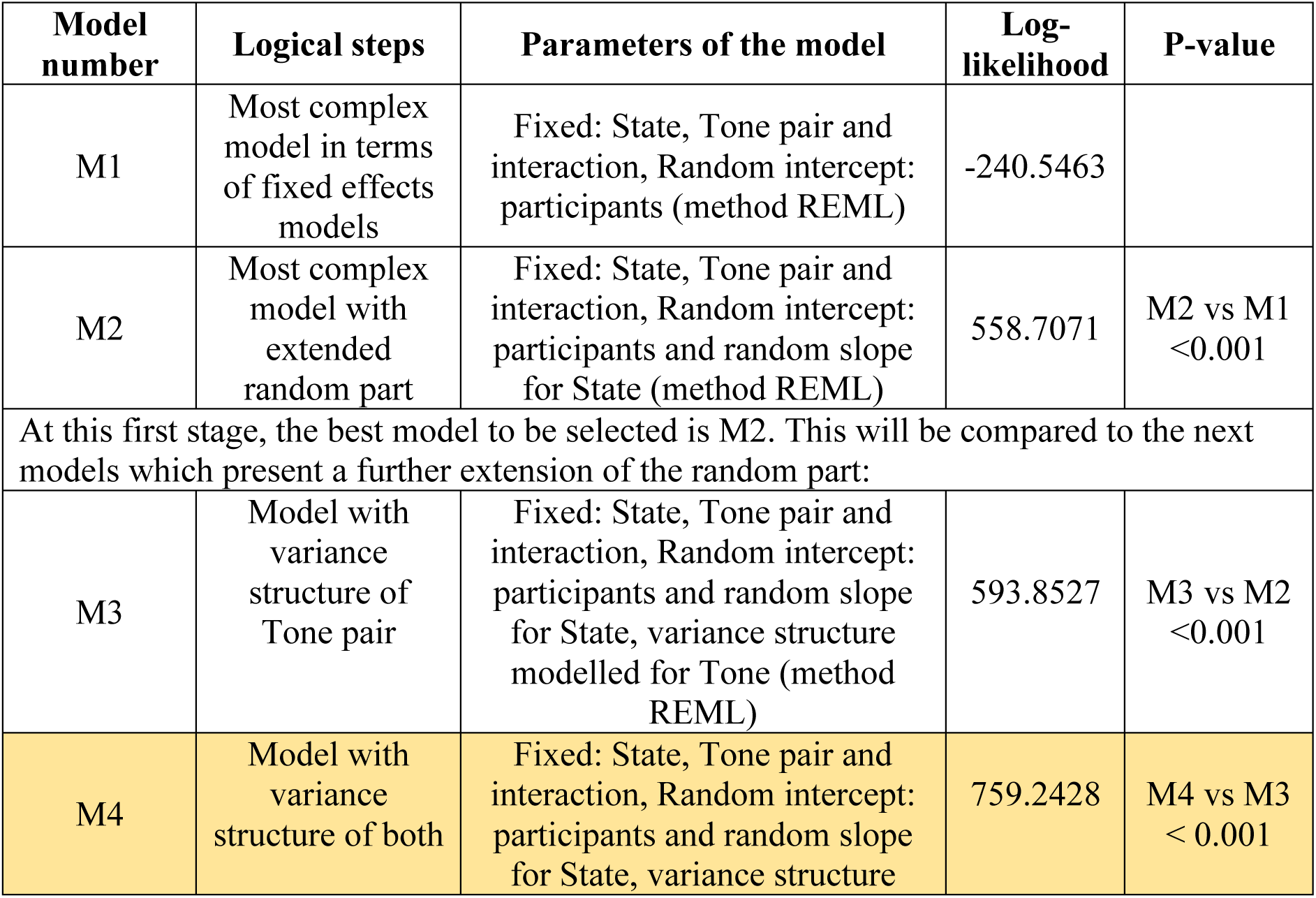
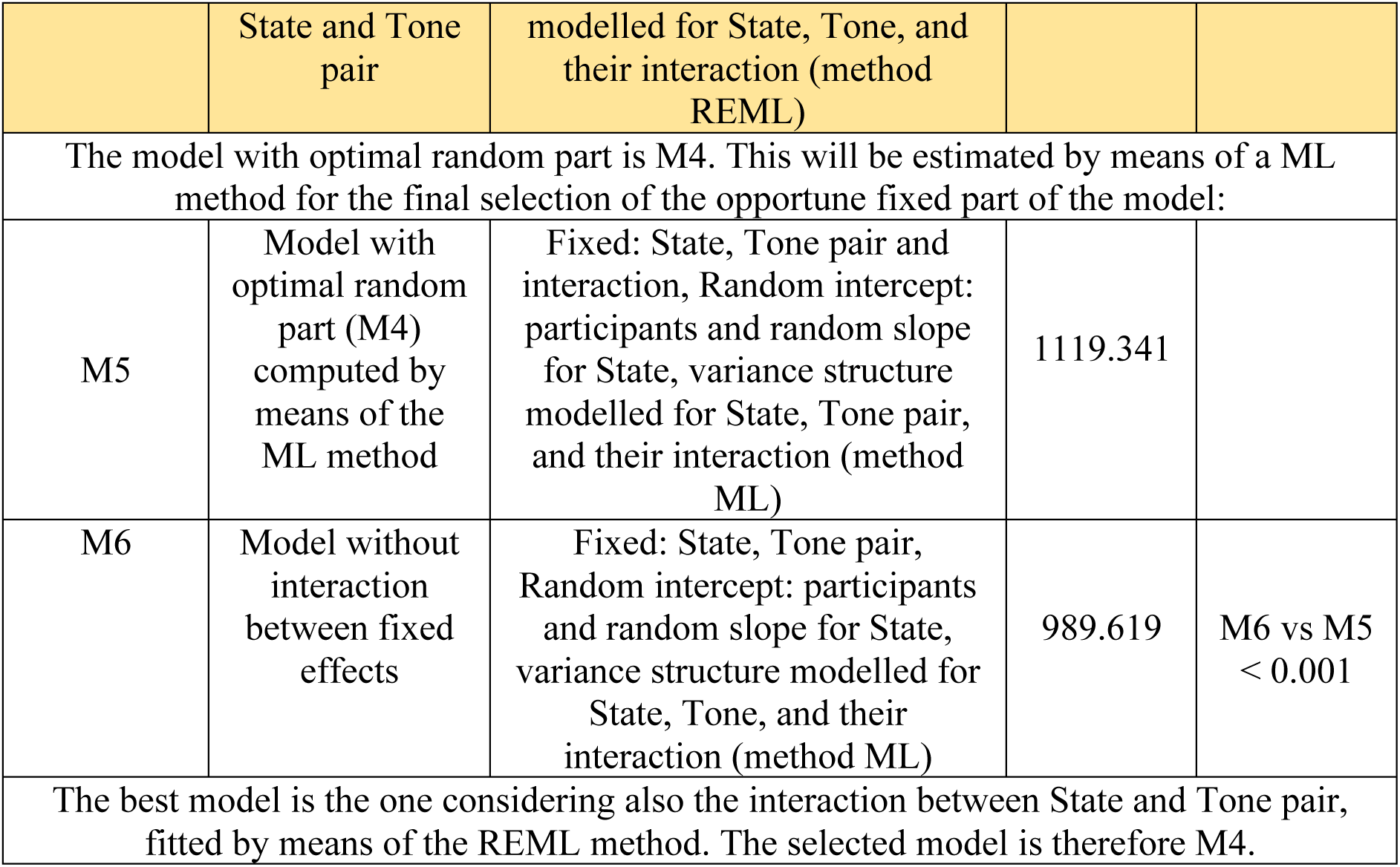
Linear mixed-effects models for the early time window.

**Table 5:** Linear mixed-effects models for the late time window.

| Model number | Logical steps | Parameters of the model | Log-likelihood | P-value |
| --- | --- | --- | --- | --- |
| M1 | Most complex model in terms of fixed effects models | Fixed: State, Tone pair and interaction, Random intercept: participants (method REML) | -918.9524 |  |
| M2 | Most complex model with extended random part | Fixed: State, Tone pair and interaction, Random intercept: participants and random slope for State (method REML) | -84.8070 | M2 vs M1 <0.001 |
| At this first stage, the best model to be selected is M2. This will be compared to the next models which present a further extension of the random part: |  |  |  |  |
| M3 | Model with variance structure of Tone pair | Fixed: State, Tone pair and interaction, Random intercept: participants and random slope for State, variance structure modelled for Tone (method REML) | -60.93222 | M3 vs M2 <0.001 |
| M4 | Model with variance structure of both State and Tone pair | Fixed: State, Tone pair and interaction, Random intercept: participants and random slope for State, variance structure modelled for State, Tone, and their interaction (method REML) | 85.89841 | M4 vs M3 < 0.001 |
| The model with optimal random part is M4. This will be estimated by means of a ML method for the final selection of the opportune fixed part of the model: |  |  |  |  |
| M5 | Model with optimal random part (M4) computed by means of the ML method | Fixed: State, Tone pair and interaction, Random intercept: participants and random slope for State, variance structure modelled for State, Tone pair, and their interaction (method ML) | 426.7567 |  |
| M6 | Model without interaction between fixed effects | Fixed: State, Tone pair, Random intercept: participants and random slope for State, variance structure modelled for State, Tone, and their interaction (method ML) | 314.8385 | M6 vs M5 < 0.001 |
| The best model is the one considering also the interaction between State and Tone pair, fitted by means of the REML method. The selected model is therefore M4. |  |  |  |  |

To disentangle and characterize possible differences in auditory processing between time windows, further analyses were conducted for early and late time windows, separately. In the early time window, linear mixed-effects modelling analysis (Table 4) revealed a main effect of *state* (F(3,5005) = 76.58, p < 0.0001, *η*^2^ = 0.04 ; Fig. 3b), a main effect of *tone-pair* (F(35, 5005) = 42.35, p < 0.0001, *η*^2^ = 0.23; Fig. 3) and an interaction between *state* and *tone-pair* (F(105, 5005) = 2.58, p < 0.0001, *η*^2^ = 0.05). Planned comparisons uncovered that the Tone Similarity Index was greater in wakefulness (r = 0.57 ± 0.17) in comparison to all sleep stages (N2: r = 0.47 ± 0.24, F(1,2485) = 5.78, p = 0.016, *η*^2^ = 0.002; N3: r = 0.22 ± 0.25, F(1,2485) = 119.2, p < 0.0001, *η*^2^ = 0.05; REM: r = 0.41 ± 0.26, F(1,2485) = 21.1, p < 0.0001, *η*^2^ = 0.008; Fig. 3b and 4a-c), indicating greater similarities between neural responses to tone-pairs in wakefulness versus sleep. In addition, during sleep, Tone Similarity Index was greater in N2 in comparison to N3 (F(1,2485) = 1253.9, p < 0.0001, *η*^2^ = 0.34; Fig. 3b and 4) and REM (F(1,2502) = 36.3, p < 0.0001, *η*^2^ = 0.19; Fig. 3b and 4f), and greater in REM in comparison to N3 (F(1,2441) = 64.9, p < 0.0001, *η*^2^ = 0.03; Fig. 3b and 4e), indicating that neural similarities between tone-pairs depended on sleep stage. To further examine how tone frequency interacts with the conscious state we conducted planned comparisons between pairs of states and tone-pairs. We found *state* and *tone-pair* interactions, between wakefulness and all sleep stages (**N2**: F(35, 2485) = 2.16, p < 0.0001, *η*^2^ = 0.03; **N3** F(35, 2485) = 2.03, p = 0.0003, *η*^2^ = 0.03; **REM**: F(35, 2485) = 1.92, p = 0.001, *η*^2^ = 0.03; Fig.3b and 4a-c), reflecting significant differences in tone-pairs similarities between states in low frequencies and no change in high frequencies (Fig. 4a-c). In addition, within sleep there were interactions between N3 and N2 (F(35, 2485) = 1.67, p = 0.008, *η*^2^ = 0.03) and between N3 and REM (F(35, 2441) = 2.53, p < 0.0001, *η*^2^ = 0.03), reflecting greater differences in tone-pairs similarities between states in high versus low frequencies (Fig. 4d-f). These interactions suggest a stimulus-dependent modulation of neural similarities between wakefulness and sleep as well as between sleep stages for the early time window.

In the late time window, a different pattern emerged. Linear mixed-effects modelling analysis (Table 5) revealed a main effect of *state* (F(3,5005) = 30.91, p < 0.0001, *η*^2^ = 0.02; Fig. 3c), a main effect of *tone-pair* (F(35, 5005) = 15.72, p < 0.0001, *η*^2^ = 0.10; Fig. 3c), and an interaction between *state* and *tone-pair* (F(105, 5005) = 2.25, p < 0.0001, *η*^2^ = 0.05; Fig. 3c and 4), similar to the early time window. However, in contrast to the early time window, Tone Similarity Index was greater in N2 (r = 0.50 ± 0.26) than in wakefulness (r = 0.25 ± 0.21, F(1,2455) = 46.6, p < 0.0001, *η*^2^ = 0.02; Fig. 3c), N3 (r = 0.30 ± 0.29, F(1,2451) = 175.8, p < 0.0001, *η*^2^ = 0.07), and REM sleep (r = 0.28 ± 0.27, F(1,2462) = 66.5, p < 0.0001, *η*^2^ = 0.03; Fig. 3c and 4), while no other differences were evident between wakefulness and sleep nor between sleep stages (all F’s < 5.25, p > 0.05 ; Fig. 3c and 4). Moreover, an interaction between *states* and *tone-pair* was observed between N2 and wakefulness (F(35, 2455) = 4.48, p < 0.0001, *η*^2^ = 0.02; Fig. 3c and 4a), and N2 and N3 (F(35, 2451) = 3.67, p < 0.0001, *η*^2^ = 0.05; Fig. 3c and 4d), reflecting greater differences in tone-pairs similarities between states in high versus low frequencies (Fig. 4a,d). In addition, *state* and *tone-pair* interactions between REM and wakefulness (F(35, 2439) = 2.94, p < 0.0001, *η*^2^ = 0.04; Fig. 3c and 4c), and REM and N3 (F(35, 2435) = 2.76, p < 0.0001, *η*^2^ = 0.04; Fig. 3c and 4f) were also found, indicating that even when the mean Tone Similarity Index is largely the same between states (i.e., no main effects of state were found between wakefulness, REM and N3), the relation between neural responses to tones is state-dependent. These findings demonstrate a differential modulation of tone-pairs neural similarities by conscious state and further support the hypothesis of stimulus-dependent auditory processing across the sleep-wake cycle.

To further understand how auditory processing changes between the early and late time windows, we compared the Tone Similarity Index between the two windows separately for each conscious state. This analysis revealed opposite trajectories between wakefulness and NREM sleep. During wakefulness Tone Similarity Index was greater in the early compared with the late time window (F(1,2485) = 130.38, p < 0.0001, *η*^2^ = 0.05; Fig. 3), while in NREM sleep Tone Similarity Index was greater in the late compared with the early time window, for both N2 (F(1,2485) = 21.23, p < 0.0001, *η*^2^ = 0.008; Fig. 3) and N3 sleep (F(1,2485) = 60.67, p < 0.0001, *η*^2^ = 0.02; Fig. 3). No difference was found in REM sleep between early and late time windows (F(1,2485) = 1.59, p = 0.21, *η*^2^ <0.001; Fig. 3), although Fig. 3a shows some fluctuations of similarity magnitude during REM sleep as well. In addition, an interaction between *time window* and *tone-pairs* was observed in wakefulness (F(35, 2485) = 3.37, p < 0.0001, *η*^2^ = 0.05) and REM sleep (F(35, 2485) = 1.96, p = 0.0007, *η*^2^ = 0.03), but not in NREM sleep, suggesting within state stimuli-dependent modulation of neural similarities across time. Overall, these findings provide additional evidence for uneven reshaping of tone-pairs neural similarities between wakefulness and sleep, as well as between sleep stages along processing time.

### The patterns of auditory neural similarities are state-, time- and stimulus-dependent

Next, we investigated how the patterns of neural similarities between tone-pairs are modulated across the sleep-wake cycle using *Representational Similarity Analysis* (RSA) (Kriegeskorte et al., 2008). Tone Similarity Index values for all pairwise combinations of the nine tone frequencies were reorganised into *Representational Similarity matrices* (RSMs), generated by arranging the tone frequencies along the rows and columns of the RSM. We obtained 9 by 9 matrices, in which each cell of the matrix indicates the neural similarity between tone-pairs as measured by the Tone Similarity Index (Fig. 5 a-d). Using RSA, the RSMs were compared between conscious states in each time window separately. First, we examined the group-averaged RSMs (see methods). In the early time window, the RSM of each state was correlated with the RSM of all other conscious states (*W-N2* r = 0.90, *W-N3* r = 0.80, *W-REM* r = 0.86, N2-N3 r = 0.72, *N2-REM* r = 0.96, *N3-REM* r = 0.65, all p’s < 0.05 FDR corrected for multiple comparisons; Fig. 5a-d). Notably, the RSM of N3 had the lowest correlation values with the RSMs of the other conscious states. In the late time windows, similarly to the early time window, RSMs were correlated to each other between all pairs of conscious states (*W-N2* r = 0.63, *W-N3* r = 0.66, *W-REM* r = 0.57, N2-N3 r = 0.78, *N2-REM* r = 0.93, *N3-REM* r = 0.74, all p’s < 0.05 FDR corrected for multiple comparisons; Fig. 5a-d). Yet, unlike in the early time window, here wakefulness RSM showed lower correlation values with all sleep stages. These findings imply a relatively preserved neural similarity patterns between tone-pairs across the sleep wake-cycle.

**Fig. 5:**
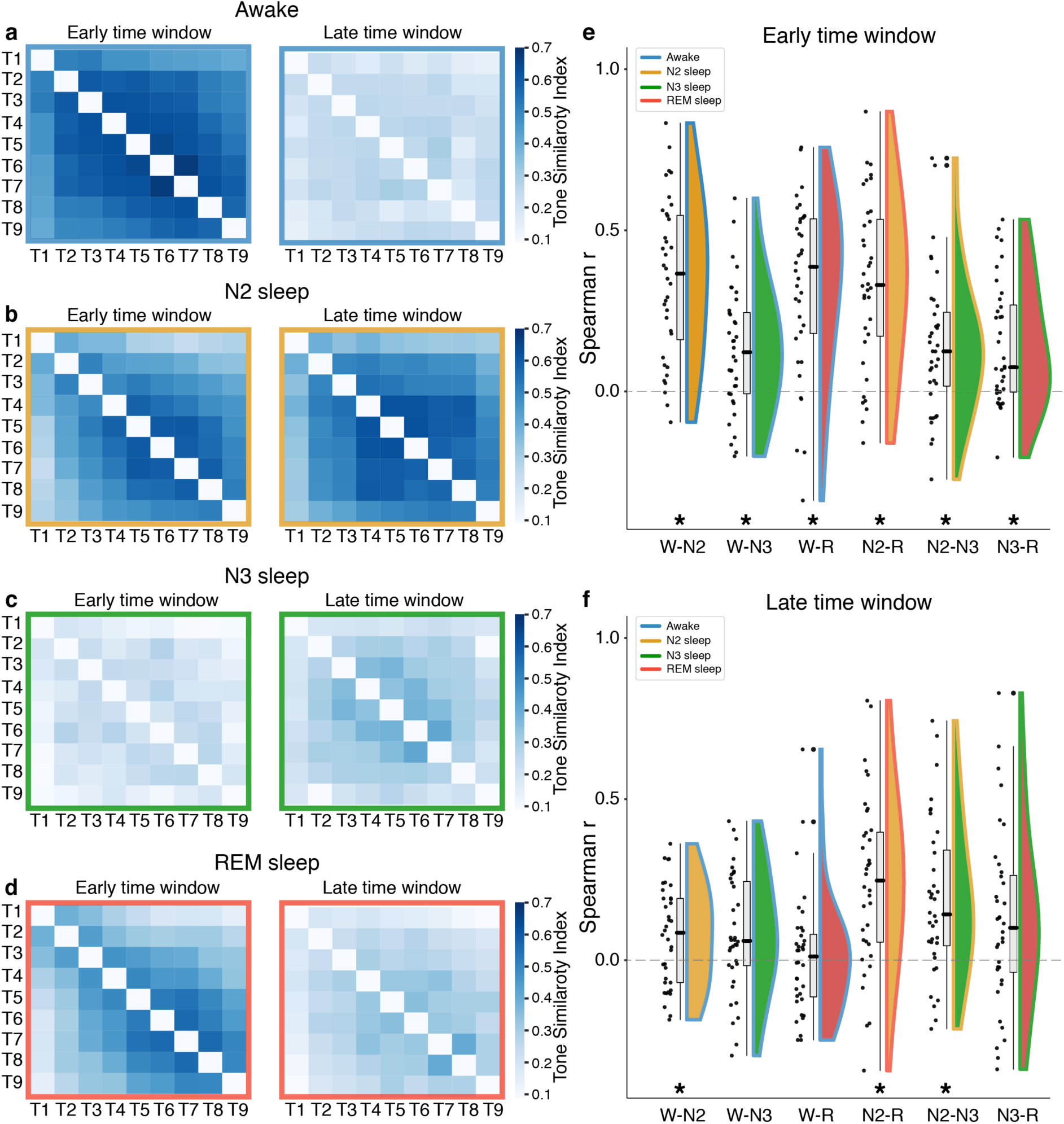
Similarity pattern between auditory responses is state-, time- and stimulus-dependent. Representational Similarity Matrices (RSM) averaged across participants during **a)** wakefulness (blue outline), **b)** N2 sleep (yellow outline), **c)** N3 sleep (green outline) and **d)** REM sleep (red outline), averaged across the early time window (left), and late time window (right). **E-f)** Individual RSMs correlations between conscious states in the **e)** early time window, where some degree of similarity was found between all conscious states, and **f)** late time window, where some degree of similarities was found only between N2 and the other conscious states. Each dot represents a participant; flat violin plots show the distribution; the boxplot mid-line denotes the median; the rectangle denotes the interquartile range (25th to the 75th percentiles). * denotes a reliable difference from zero.

To further investigate the contribution of each participant to the observed group similarity patterns and to be able to generalize such results to other samples of participants, we conducted an RSA by computing separate RSMs for each participant at each state (see methods). In the early time window, all conscious states showed some degree of RSMs similarity to each other, as in the group-averaged RSMs (***W-N2***: r = 0.35 ± 0.24, CI = [0.24, 0.47]; ***W-N3***: r = 0.13 ± 0.19, CI = [0.041, 0.22]; ***W-REM:*** r = 0.33 ± 0.26, CI = [0.20, 0.45]; ***N2-N3***: r = 0.14 ± 0.22, CI = [0.44, 0. 253]; ***N2-REM***: r = 0.34 ± 0.26, CI = [0.22, 0.46]; ***N3-REM:*** r = 0.15 ± 0.19, CI = [0.06, 0.24]; Fig. 5e). However, in the late time window, only the RSM of N2 correlated with all other states (***W-N2***: r = 0.08 ± 0.15, CI = [0.03, 0.144], ***N2-N3***: r = 0.19 ± 0.24, CI = [0.08, 0.30], ***N2-REM***: r = 0.24 ± 0.26, CI = [0.11, 0.36]), while the RSMs of wakefulness, N3 and REM sleep exibited non-reliable correlations implying a different organisation. (***W-REM***: r = 0.01 ± 0.19, CI = [-0.07, 0.11]; ***W-N3***: r = 0.08 ± 0.18, CI = [-0.003, 0.17]; ***N3-REM***: r = 0.12 ± 0.27, CI = [-0.005, 0.25]; Fig. 5f). These findings show that although wakefulness, N3 and REM sleep do not differ in Tone Similarity Index values (Fig. 3c), they present different similarity *pattern* (Fig. 5), as alluded by the state-tone interactions between these states in the similarity *magnitude* analysis (Fig. 3c). In addition, comparison of RSMs between early and late time windows in each state revealed highest correlations in N2 (r = 0.40 ± 0.25, CI = [0.29, 0.50]), then in REM (r = 0.26 ± 0.25, CI = [0.16, 0.36]), wakefulness (r = 0.19 ± 0.22 CI = [0.10, 0.28]), and lastly during N3 sleep (r = 0.13 ± 0.19, CI = [0.05, 0.21]), highlighting greater differences in similarity pattern across time for N3 sleep and more preserved pattern in N2 sleep. To further illustrate the relationships between conscious states, time windows, and tone-pairs we applied hierarchical clustering analysis. Dendrograms from this analysis provide an intuitive representation of the Tone Similarity Index magnitude (Fig. 6a-b) and patterns (Fig. 6c-l). For example, the dendrograms clearly demonstrate that N3 sleep in early time window is the most dissimilar from the other states in terms of neural similarity magnitude (Fig. 6a) and pattern (Fig. 6c), suggesting that slow wave sleep is characterised by a markedly different sensory organisation from all other states. In addition, it is also evident that N2 and REM sleep maintain a high resemblance in similarity patterns in in both time windows (Fig. 6c-d), despite their dissimilarity in magnitude, particularly for the late time window (Fig. 6b).

**Fig. 6:**
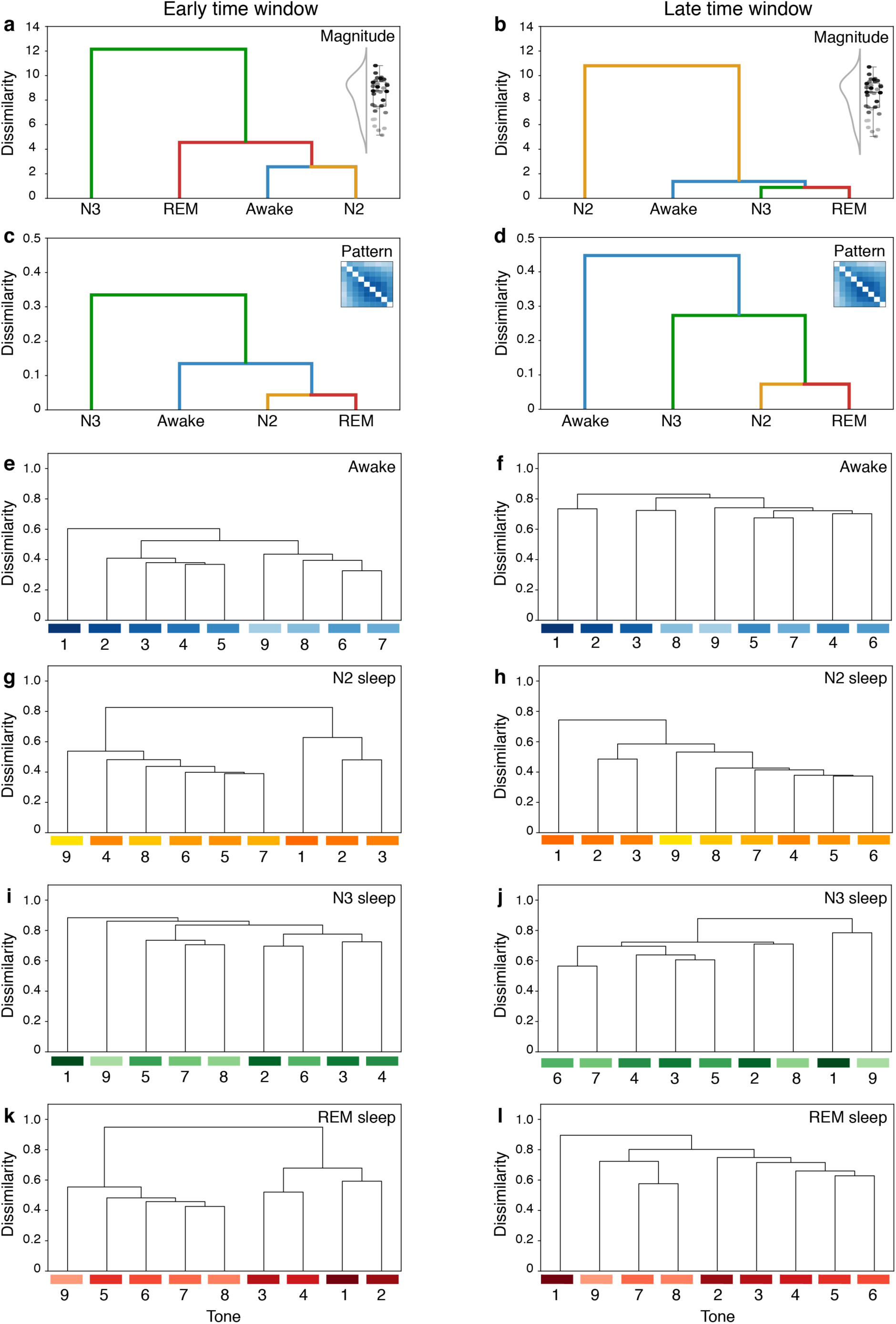
Hierarchical clustering analysis for Similarity magnitude and pattern reveals reshaping of the auditory map a-d) Hierarchical clustering analysis between conscious states in the early (left) and late (right) time windows**. a-b)** similarity magnitude across tone and **c-d)** similarity pattern across tones. **e-f)** Hierarchical clustering analysis for similarity pattern between tones in wakefulness (blue palette), **g-h)** N2 sleep (yellow palette), **i-j)** N3 sleep (green palette) and **k-l)** REM sleep (red palette). T1 =650Hz, T2 =845Hz, T3 =1098Hz, T4 =1428Hz, T5 =1856Hz, T6 =2413Hz, T7 =3137Hz, T8 =4079Hz and T9 =5302Hz.

### Probing the neural auditory similarities to auditory physical, physiological and perceptual models

Last, we tested how well the neural similarity patterns in wakefulness, NREM and REM sleep could be explained by competing conceptual models of physical, physiological and perceptual auditory organizations. The first model is the *Frequency Difference Model*, which is based on the physical distance in Hertz between tone frequencies. In this model, the relations between tone-pairs were calculated by subtracting the lower tone frequency from the higher tone frequency (Fig. 7a). The second model is the *Greenwood Model*, which was developed based on studies mathematically defining the links between the anatomic location of the inner ear hair cells in the cochlea and the tone frequencies at which they are stimulated (Greenwood, 1990). Here, the relations between tone-pairs was calculated by applying the Greenwood function, which estimates the distance in millimetres along the basilar membrane between locations that are maximally excited by each frequency (Fig. 7b, See methods), representing “physiological distances”. The Third model is the *Mel Model*, obtained from the Mel scale, which indicates how pitch perception evolves as a function of sound frequencies in a non-linear manner (Micheyl et al., 2012; Moore, 2003; Stevens et al., 1937). In the perceptual model, the relations between tone-pairs were calculated by dividing the Mel value corresponding to the higher tone frequency by Mel value corresponding to the lower tone frequency (Fig. 7c, See methods). For each of the three model, the tone-pairs relations were arranged into 9 by 9 matrices as the RSMs discussed in the previous section. To evaluate each model’s capacity to predict the observed neural relation between tone-pairs, we transformed the RSMs into *Representational Dissimilarity matrices* (RDMs = 1-RSMs) and estimated, for each participant in each conscious state and time window, the correlation between the RDMs and the model matrices. We then fitted the correlation values with a linear mixed-effects model, where *model* (‘Difference’, ’Greenwood’, ‘Mel’), *state* (‘Wakefulness’, ‘N2’, ‘N3’, ‘REM’) and *time window* (‘Early’, ‘Late’) appeared as fixed effects (Table 6). We found a main effect of *model* (F(2,811) = 14.7, p < 0.0001, *η*^2^ = 0.03), a main effect of *state* (F(3,811) = 5.44, p = 0.003, *η*^2^ = 0.02) and a main effect of *time window* (F(1,811) = 49.84, p < 0.0001, *η*^2^ = 0.06), the latter reflecting higher model fits for the early time window (Fig. 7). In addition, there were interactions between *model* and *state* (F(6,811) = 5.53, p < 0.0001, *η*^2^ = 0.04), *model* and *time window* (F(2,811) = 7.76, p = 0.0005, *η*^2^ = 0.02) and state and time window (F(3,811) = 47.03, p < 0.0001, *η*^2^ = 0.15).

**Fig. 7:**
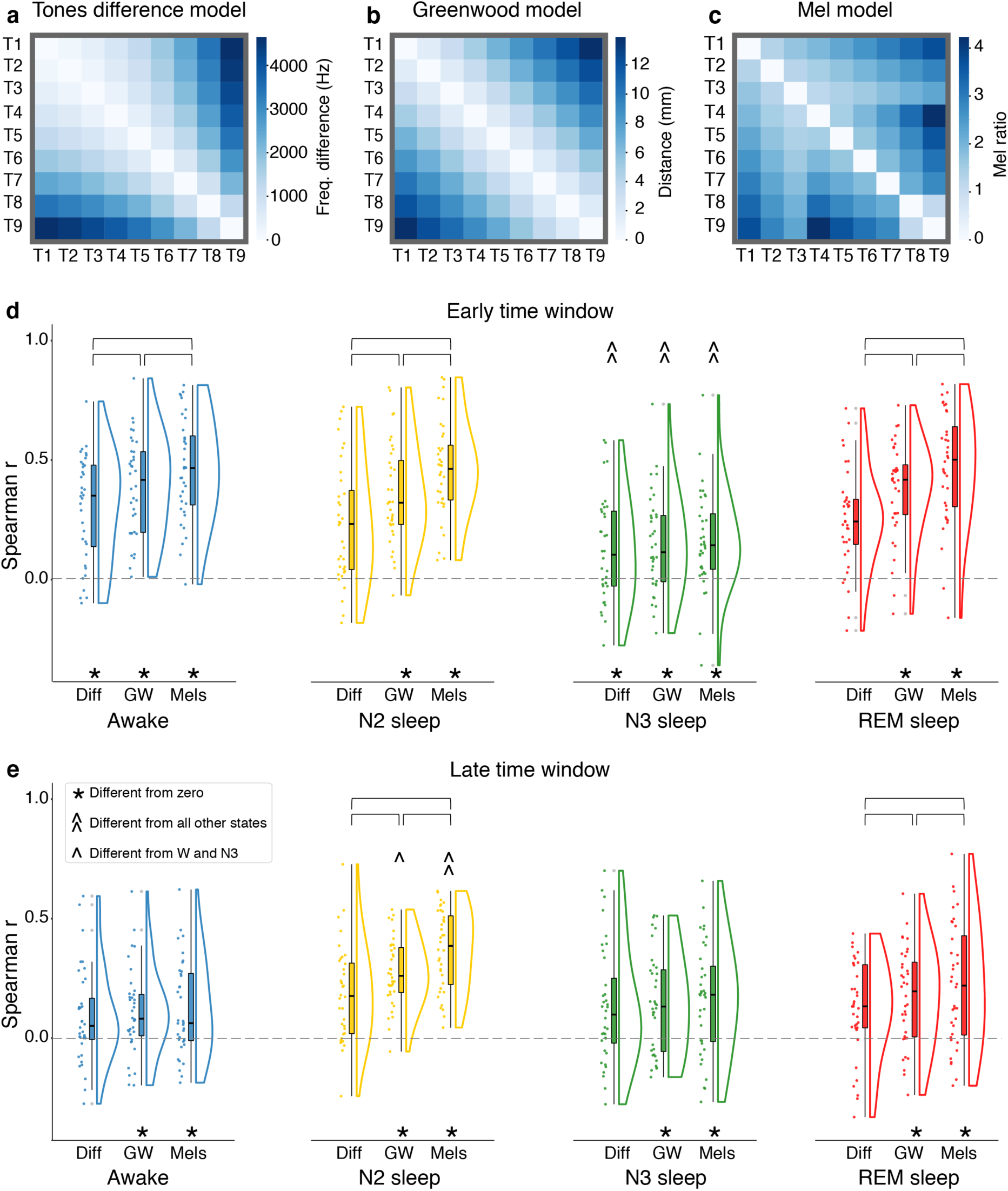
Probing the auditory neural similarities to auditory perceptual, physiological and physical models. a) The *Frequency Difference Model* reflecting the physical distance in Hz between stimuli. The value in each cell was calculated by subtracting the lower tone frequency from the higher tone frequency. **b)** The *Greenwood Model*, linking the anatomic location of the inner ear hair cells to the tone frequencies at which they are stimulated. The value in each cell was calculated by means of the Greenwood function and corresponds to the distance in millimetres along the basilar membrane between locations maximally excited by each tone frequency. **c)** The *Mel Model*, which is based on the Mel Scale relates the perceived similarity in pitch to frequency similarity while accounting for the nonlinearity of humans’ perceptual discrimination ability. Values in each cell were calculated by dividing the higher Mel value by the lower Mel value obtained for each tone-pair. Spearman’s correlations between neural similarity patterns (representation dissimilarity matrices, RDM) and auditory models show that the Mel model has higher predictive value in all conscious states, except for N3 sleep, in both **e-g)** the early time window and **h-j)** the late time window.

**Table 6:**
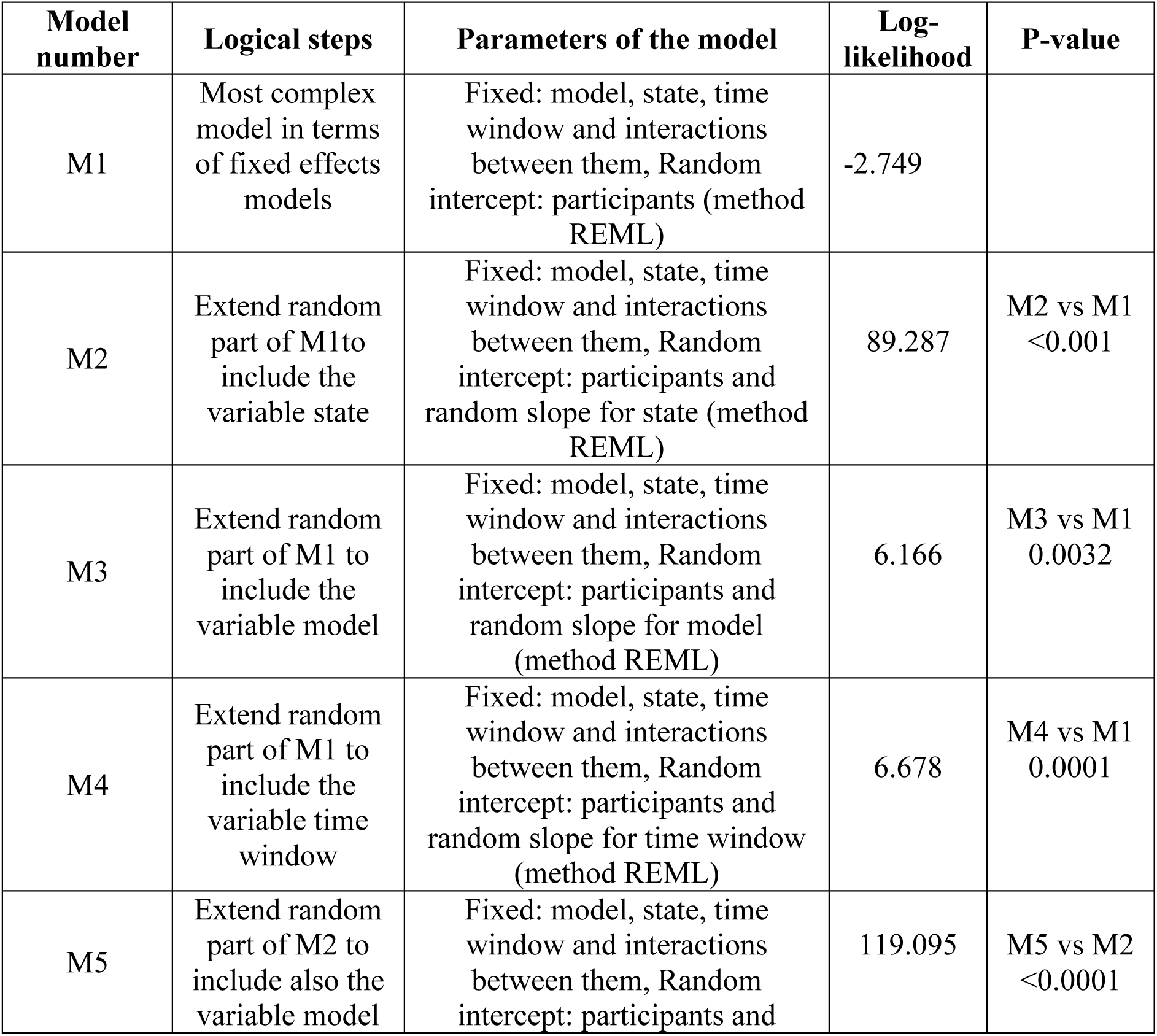
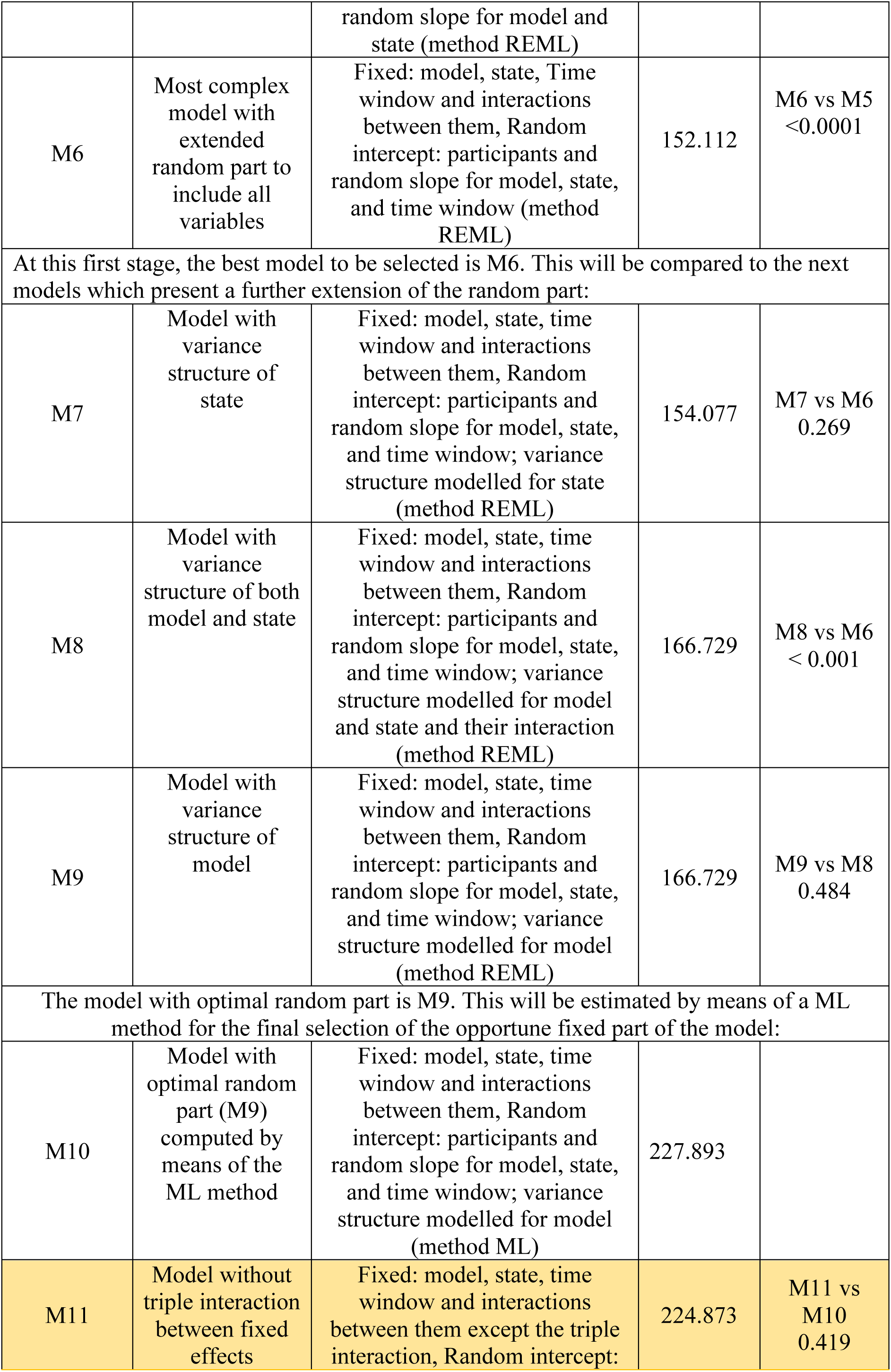
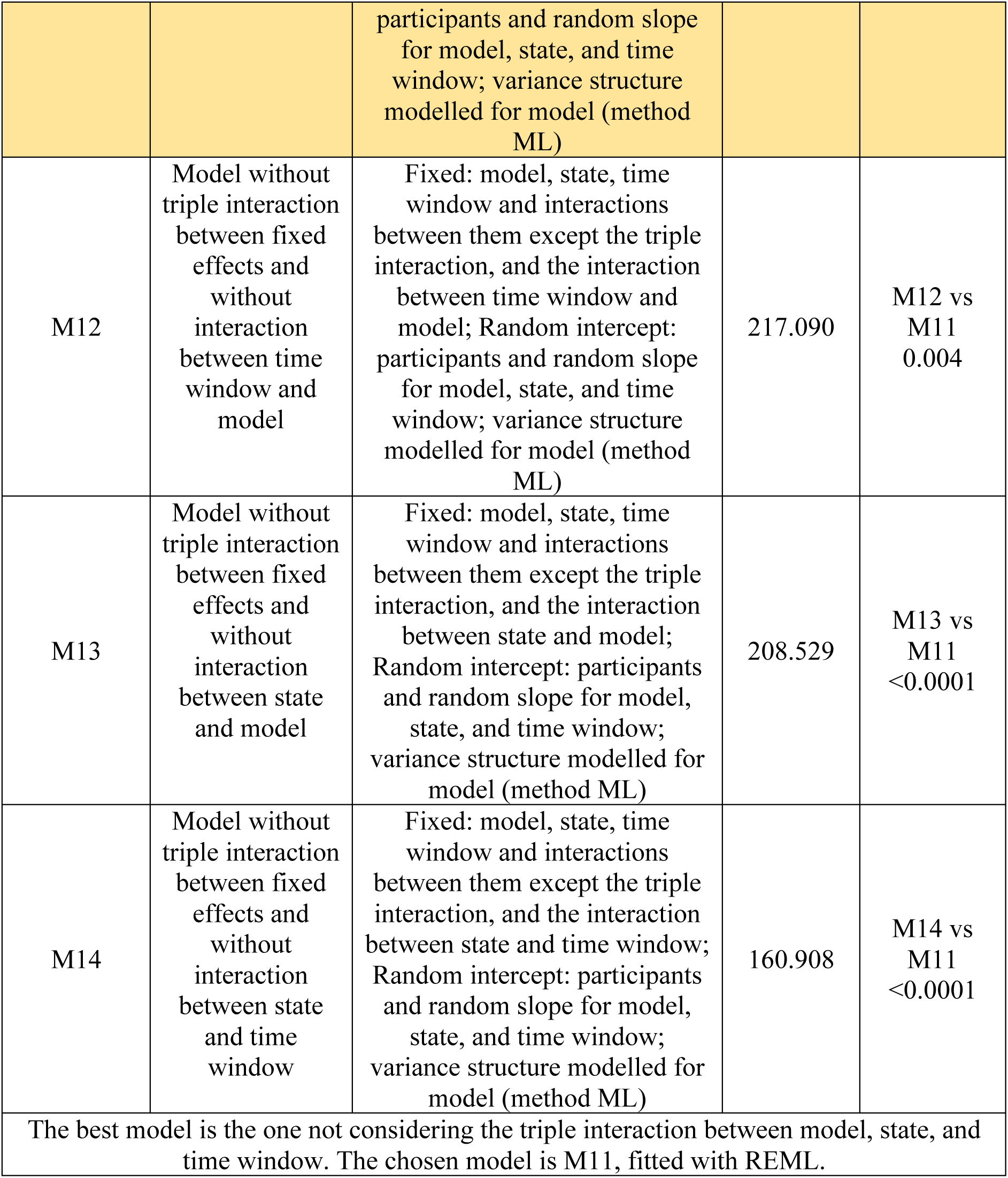
Linear mixed-effects models for auditory model analysis to assess the predictive value of the considered conceptual models.

| Model number | Logical steps | Parameters of the model | Log-likelihood | P-value |
| --- | --- | --- | --- | --- |
| M1 | Most complex model in terms of fixed effects models | Fixed: model, state, time window and interactions between them, Random intercept: participants (method REML) | -2.749 |  |
| M2 | Extend random part of M1 to include the variable state | Fixed: model, state, time window and interactions between them, Random intercept: participants and random slope for state (method REML) | 89.287 | M2 vs M1<br><0.001 |
| M3 | Extend random part of M1 to include the variable model | Fixed: model, state, time window and interactions between them, Random intercept: participants and random slope for model (method REML) | 6.166 | M3 vs M1<br>0.0032 |
| M4 | Extend random part of M1 to include the variable time window | Fixed: model, state, time window and interactions between them, Random intercept: participants and random slope for time window (method REML) | 6.678 | M4 vs M1<br>0.0001 |
| M5 | Extend random part of M2 to include also the variable model | Fixed: model, state, time window and interactions between them, Random intercept: participants and random slope for model and state (method REML) | 119.095 | M5 vs M2<br><0.0001 |
| M6 | Most complex model with extended random part to include all variables | Fixed: model, state, Time window and interactions between them, Random intercept: participants and random slope for model, state, and time window (method REML) | 152.112 | M6 vs M5<br><0.0001 |
| At this first stage, the best model to be selected is M6. This will be compared to the next models which present a further extension of the random part: |  |  |  |  |
| M7 | Model with variance structure of state | Fixed: model, state, time window and interactions between them, Random intercept: participants and random slope for model, state, and time window; variance structure modelled for state (method REML) | 154.077 | M7 vs M6<br>0.269 |
| M8 | Model with variance structure of both model and state | Fixed: model, state, time window and interactions between them, Random intercept: participants and random slope for model, state, and time window; variance structure modelled for model and state and their interaction (method REML) | 166.729 | M8 vs M6<br>< 0.001 |
| M9 | Model with variance structure of model | Fixed: model, state, time window and interactions between them, Random intercept: participants and random slope for model, state, and time window; variance structure modelled for model (method REML) | 166.729 | M9 vs M8<br>0.484 |
| The model with optimal random part is M9. This will be estimated by means of a ML method for the final selection of the opportune fixed part of the model: |  |  |  |  |
| M10 | Model with optimal random part (M9) computed by means of the ML method | Fixed: model, state, time window and interactions between them, Random intercept: participants and random slope for model, state, and time window; variance structure modelled for model (method ML) | 227.893 |  |
| M11 | Model without triple interaction between fixed effects | Fixed: model, state, time window and interactions between them except the triple interaction, Random intercept: | 224.873 | M11 vs M10<br>0.419 |

|  |  | participants and random slope for model, state, and time window; variance structure modelled for model (method ML) |  |  |
| --- | --- | --- | --- | --- |
| M12 | Model without triple interaction between fixed effects and without interaction between time window and model | Fixed: model, state, time window and interactions between them except the triple interaction, and the interaction between time window and model; Random intercept: participants and random slope for model, state, and time window; variance structure modelled for model (method ML) | 217.090 | M12 vs M11<br>0.004 |
| M13 | Model without triple interaction between fixed effects and without interaction between state and model | Fixed: model, state, time window and interactions between them except the triple interaction, and the interaction between state and model; Random intercept: participants and random slope for model, state, and time window; variance structure modelled for model (method ML) | 208.529 | M13 vs M11<br><0.0001 |
| M14 | Model without triple interaction between fixed effects and without interaction between state and time window | Fixed: model, state, time window and interactions between them except the triple interaction, and the interaction between state and time window; Random intercept: participants and random slope for model, state, and time window; variance structure modelled for model (method ML) | 160.908 | M14 vs M11<br><0.0001 |
| The best model is the one not considering the triple interaction between model, state, and time window. The chosen model is M11, fitted with REML. |  |  |  |  |

Planned comparisons revealed that in the early time window across all states the Mel model had a higher predictive value than the Greenwood (t(811) = 4.37, p < 0.0001. Cohen’s d = 0.19; Fig. 7) and Difference (t(811) = 6.52, p < 0.0001. Cohen’s d = 0.39; Fig. 7e-g) models, and that the Greenwood model had a higher predictive value than the Difference model (t(811) = 5.28, p < 0.0001. Cohen’s d = 0.20; Fig. 7e-g). In the late time window, the Mel model had a higher predictive value than the Difference model (t(811) = 3.07, p = 0.006. Cohen’s d = 0.185; Fig. 7h-j) but no reliable difference was found between the Mel and Greenwood models (t(811) = 2.32, p = 0.054. Cohen’s d = 0.10; Fig. 7h-j), nor between the Difference and Greenwood models (t(811) = 2.20, p = 0.072, Cohen’s d = 0.08; Fig. 7h-j). These findings suggest that not only in wakefulness but also during sleep, tone-pairs neural similarities organisation is better explained by the perceived similarities between tone frequencies than by the tones difference in Hz or by the ear anatomo-physiological organization.

Nonetheless, these results do not imply that the *Models* fit the auditory neural representations to the same extent in all states. Indeed, in the early time window the three models had lower predictive value for N3 in comparison to all other states (all t’s > 3.55, all p’s < 0.0023), in line with the similarity magnitude (Fig. 3) and pattern (Fig. 5) results, signalling distinctive neural auditory frequency organisation in N3. Furthermore, in the late time window, the Mel model had a higher predictive value for N2 in comparison to all other states (all t’s > 2.67, all p < 0.039, Cohen’s d > 0.25), and the Greenwood model had a higher predictive value for N2 in comparison to wakefulness (t(811) = 3.87, p = 0.007, Cohen’s d = 0.33) and N3 (t(811) = 3.68, p = 0.001, Cohen’s d = 0.26), but not in relation to REM sleep (t(811) = 2.47, p = 0.065, Cohen’s d = 0.21). No reliable differences were found between states for the Difference model in the late time window. Altogether, these findings indicate that the general rules based on perceptual similarities governing the structure of the tone-pairs neural similarity organisation during wakefulness, are also applicable to sleep, yet in a conscious-state sleep stage-dependent manner.

## Discussion

In this study, we sought to examine how conscious states shape neural sensory mapping of the auditory system in humans. We systematically examined brain activity in response to a range of pure tones during wakefulness, NREM and REM sleep and revealed that the tone-pairs neural similarities are dynamically changing in a state-, time- and stimulus-dependent manner. Specifically, the neural similarity between pure tones differed between wakefulness and sleep, as well as between sleep stages in both early and late auditory processing windows, revealing different hierarchical relationships between states across time. Furthermore, tone-pairs neural similarities were modulated by conscious state as a function of tone frequency, with some tones-pairs showing a change in similarity between states while others remain unchanged. In accordance with our hypothesis, these findings demonstrate convergent evidence of functional auditory reorganisation across the sleep-wake cycle.

Over the last century, the anatomical and functional organization of sensory systems have been vastly studied in awake and sedated animal models (Hudspeth and Logothetis, 2000; Kaas, 2008). For many years, it has been assumed that the sensory maps in the brain are resistant to changes in conscious states, at least at the level of primary sensory cortices. This assumption turned out to be inaccurate (Reimann and Niendorf, 2020), and raised the question of the stability and flexibility of sensory maps between different states of consciousness. In this study, we investigated how the functional organisation of the auditory frequency map is shaped across the sleep-wake cycle and revealed that it is highly sensitive to changes in conscious state. These findings are in line with a series of studies clearly showing that sensory maps are flexible and dynamic in wakefulness. Sensory remapping can occur following changes in context (Hayman et al., 2003), attention (Berman and Colby, 2009; Burr and Morrone, 2011; Fritz et al., 2003; Rolfs and Szinte, 2016), or learning (Banerjee et al., 2020; Bostock et al., 1991; Leutgeb et al., 2005), and it can be expressed in different forms, such as global, partial or conditional remapping (Fyhn et al., 2007; Jeffery, 2011; Rennó-Costa et al., 2010), in a range of modalities (Geva-Sagiv et al., 2016; Heed et al., 2015; Lee et al., 2003; Rolfs and Szinte, 2016). In the auditory modality, there are numerous examples for dynamic adjustments of neural responses to changes in context, in wakefulness (Blake and Merzenich, 2002; Fritz et al., 2003; Garrido et al., 2013; Regev et al., 2020), and under anaesthesia (Gourévitch et al., 2009). In addition, there is evidence for reorganisation of tonotopic maps following learning (Gonzalez-Lima and Agudo, 1990; Scheich, 1991), attention to a specific tone (Dick et al., 2017), or auditory deprivation and rehabilitation (Thai-Van et al., 2010).

Here, we contribute to these findings by uncovering commonalities and differences in neural auditory frequency organisation between wakefulness and different sleep stages. First, the neural auditory frequency map assessed by means of neural similarities between tone-pairs changed along auditory processing time and between conscious states. Specifically, *Tone Similarity Index* (see methods), decreased in wakefulness, increased during NREM sleep and remained on average unchanged during REM sleep across the processing time. Moreover, within each processing time window, a different set of hierarchical relationships emerged between states. During the early processing window, Tone Similarity Index magnitude was higher in wakefulness in comparison to all sleep stages, and within sleep, it was highest in N2, then in REM and lastly N3 sleep. This hierarchy was altered in the late processing window, where Tone Similarity Index magnitude in N2 was higher than in all other states, and did not differ between wakefulness, N3 and REM sleep. Together, these findings reveal a dynamic reorganisation of the auditory neural frequency map across the sleep-wake cycle (Fig. 3). Second, the neural auditory frequency map structure as assessed by means of RSA, showed some degree of pattern preservation across all states, together with clear differences between them. Specifically, interactions between tones-pairs and states were observed between all states, with the exception of N2 and REM. Indeed, in both time windows, N2 and REM sleep presented a relatively preserved auditory frequency map organisation, despite the observed changes in the mean Tone Similarity Index magnitude between the two states. In other words, tone-pairs that are relatively similar in N2 are also relatively similar in REM sleep and vice versa, irrespective of Tone Similarity Index magnitudes. The opposite pattern - reorganisation of map assembly while maintaining magnitudes - was evident between wakefulness, N3 and REM sleep in the late processing window. Collectively, these findings uncover different forms of reorganisation across the sleep-wake cycle, including modulation of Tone Similarity Index magnitudes without a change in pattern and a modulation of Tone Similarity Index pattern without a change in magnitudes.

The observed functional reorganisation across the sleep-wake cycle dovetails nicely with several studies showing that the degree of brain modulation in sleep is sensitive to the properties of the stimulus (Castro-alamancos, 2004; Issa and Wang, 2011; Lustenberger et al., 2018; Portas et al., 2000; Sharon and Nir, 2018; Tlumak et al., 2012). For example, stimulus presentation rate has a bidirectional effect on steady-state evoked responses, where low-frequency stimulation elicits stronger responses during sleep, while high-frequency stimulation elicits stronger activation during wakefulness (Castro-alamancos, 2004; Lustenberger et al., 2018; Sharon and Nir, 2018; Tlumak et al., 2012). In addition, stimulus intensity could impact neural activity in an unequal manner, such that firing rate to quiet but not loud sounds is reduced during sleep (Issa and Wang, 2011), and the change in amplitude of steady state evoked potentials with increased tone intensity is smaller in sleep than in wakefulness (Lindens et al., 1985). Also, stimulus types are weighted differently by conscious states, with sleep showing selective enhancement of BOLD responses over wakefulness for one’s own name in comparison to beeps (Portas et al., 2000), and divergent oscillatory responses for familiar versus unfamiliar stimuli in wakefulness and different sleep stages (Blume et al., 2017, 2018). In addition, at the behavioural level, increased arousal probability from sleep was found for one’s own infant cry (Formby, 1967), as well as for low (400Hz and 520Hz) versus high (3000Hz) tone frequency (Bruck et al., 2009). These stimulus-dependent responses can potentially explain some of the discrepancies between sleep studies showing enhanced (Colrain and Campbell, 2007; Hall and Borbely, 1970; Nicholas et al., 2006; Yang and Wu, 2007), reduced (Czisch et al., 2002, 2004; Murata and Kameda, 1963) or preserved activity between sleep and wakefulness (Edeline et al., 2001; Issa and Wang, 2008; Nir et al., 2015; Peña et al., 1999). Furthermore, while responses across a neural population may show one pattern, a detailed investigation of single neurons has revealed heterogenous responses and different degrees of attenuation (Edeline et al., 2001; Issa and Wang, 2008; Nir et al., 2015; Peña et al., 1999; Sela et al., 2016, 2020). Specific properties at the neuronal level such as latency, selectivity or receptive field size (Edeline et al., 2001; Sela et al., 2020) may explain part of the patterns that are seen at the cortical level. The evidence gathered at the neuronal, population and cortical levels call for a comprehensive examination of a range of stimulus properties in order to unveil the interplay between conscious state and sensory mapping. Here, we directly address this need by systematically characterizing neural dynamics between a range of tone frequencies, and uncover diverse stimulus-state interactions between wakefulness and sleep and between sleep stages.

Auditory theory has advanced in many fronts using physical, physiological, neural and psychophysics evidence to build an explanatory corpus of how animals and humans hear (de Boer, 1980, 1984, 1991; Moore, 2003; Schnupp et al., 2011; Wever, 1949). Among the many models that can be applied to characterise the structure of the neural auditory frequency map in each conscious state (Meddis et al., 2010), we applied a set of models based on the perceptual (Micheyl et al., 2012; Moore, 2003; Stevens et al., 1937), physiological (Greenwood, 1990), and physical relation between tones frequencies. We found that the neural auditory frequency map structure was better described by the perceptual relation between tones which reflect pitch discriminability than by differences between tones’ physical properties, not only in wakefulness but also in N2 and REM sleep. It is well accepted that in wakefulness pitch perception is based more on the neural representation of sound at the output of the auditory periphery than on the physical properties of sound as it enters the ear (Moore, 2003; Yost, 2009). During sleep, modulation of auditory responses is evident at the cortical (Blume et al., 2017, 2018; Czisch et al., 2002; Dang-Vu et al., 2010; Portas et al., 2000; Schabus et al., 2012; Strauss et al., 2015; Wilf et al., 2016), and sub-cortical levels of the auditory pathway including the thalamus (Edeline et al., 2000; Hall and Borbely, 1970), the inferior colliculus (Morales-cobas et al., 1995), the lateral superior olive (Pedemonte et al., 1994), and the cochlea (Froehlich et al., 1993; Irvine and Webster, 1972; Velluti et al., 1989). Yet, despite the vast modulation of auditory processing in sleep, we show here that the neural auditory frequency map preserves an organisation that follows perceptual similarities during both NREM and REM sleep, with the exception of N3. These findings imply some level of functional auditory organisation stability between conscious states.

In addition, the comparison to auditory models also uncovered the flexibility of the functional auditory organisation between conscious states. In details, during the early processing window, all three models inadequately described the auditory frequency map organisation in N3 in comparison to the other sleep stages and wakefulness (Fig. 7d). These findings are in accordance with our results from the similarity magnitude and pattern analyses, which pointed towards a markedly different sensory organization in deep sleep: N3 displayed both lower Tone Similarity Index magnitudes (Fig. 3 and 6) and a more dissimilar pattern of Tone Similarity Index in comparison to all other states (Fig. 5 and 6). The altered auditory map structure presented in N3 might be explained by findings from whole-brain fMRI studies showing a global decrease in effective interactions and breakdown of inter-modular connectivity during deep sleep (Horovitz et al., 2009; Jobst et al., 2017; Tagliazucchi et al., 2013; Tarun et al., 2021). Furthermore, the distinct auditory neural organisation during N3, may help to explain the decline in sensory and cognitive processes that is typically found during deep sleep (Andrillon and Kouider, 2020; Andrillon et al., 2016, 2017; Hennevin et al., 2007; Legendre et al., 2019; Peigneux et al., 2001). In the late processing window, a different pattern was observed: the auditory frequency map organisation was better explained by the perceptual and physiological models in N2, in comparison to wakefulness and other sleep stages. A possible interpretation for the increased predictive value as well as for the increased similarity magnitude in N2 in the late processing window might be related to delayed processing occurring during the transition from wakefulness to sleep (Atienza et al., 2001; Bastuji and García-Larrea, 1999; Canales-Johnson et al., 2019; Hennevin et al., 2007; Kouider et al., 2014; Noreika et al., 2020; Strauss and Dehaene, 2019; Strauss et al., 2015; Velluti, 1997). However, it has to be noted that our results seem to point towards a prolonged sensory processing (Fig. 3) in addition to delayed one in N2 (Fig. 2), and in order to disentangle between the mechanism of delayed and prolonged sensory processing in sleep further studies are needed.

Although N2 and N3 seem to be characterized by different auditory map organizations, we did not observe any differences in ERPs between the two states, unlike other studies showing increased ERP amplitude with sleep depth (Nielsen-Bohlman et al., 1991; Picton et al., 2003; Winter et al., 1995; Yang and Wu, 2007). This discrepancy may be driven by features that are specific to each sleep stage such as K-complexes and spindles during N2 and slow waves during N3 (Iber et al., 2007) and by their non-uninform distribution across the cortical surface (Geva-Sagiv and Nir, 2019; Siclari and Tononi, 2017). The magnitude of these sleep markers is much larger than that of the ERP’s and can strongly influence brain responses to external stimuli (Antony and Paller, 2017; Czisch et al., 2009; Dang-vu et al., 2011; Lustenberger et al., 2018; Schabus et al., 2012). Furthermore, this influence could even have a larger impact for underpowered studies. Here, stimuli were presented across a full-night of sleep accumulating thousands of repetitions per state (Table 1). The large number of trials enables averaging out ongoing brain activity and minimizing the influence of sleep stage specific features on the evoked responses while lending statistical strength to each comparison. Thus, if indeed previously observed differences between N2 and N3 are due to ongoing brain activity, the large number of stimuli employed here might explain the lack of differences in ERPs between N2 and N3 sleep in this study.

How sleep stages differ between them and from the wake state has been addressed in spontaneous brain activity (Brodbeck et al., 2012; Iber et al., 2007; Jagannathan et al., 2018; Jobst et al., 2017; Nir et al., 2015; Tagliazucchi et al., 2013), sensory processing (Andrillon and Kouider, 2020; Hennevin et al., 2007; Velluti, 1997), and cognitive dynamics (Andrillon et al., 2016, 2017, Arzi et al., 2012, 2014; Koroma et al., 2020; Strauss et al., 2015). The relationship between the degree of processing in different conscious states and the extent of sensory remapping require further investigation under the umbrella of cognitive neuroscience of unconsciousness (Chennu and Bekinschtein, 2012) and a solid theoretically and methodologically neuroscience framework (Frégnac and Bathellier, 2015; Guest and Martin, 2020; Kriegeskorte and Douglas, 2018; Lopes da Silva, 2013; Schreiner and Winer, 2007). The development of maps in neuroscience enhances the understanding of normal neural organization, its modification by pathology, and modulations by experience and context. These maps, like those charted here, serve the computational principles that govern sensory processing and the generation of perception (Schreiner and Winer, 2007) even in unconscious states (Goupil and Bekinschtein, 2012). Sleep plays, as a theoretical tool and experimental model, a key role in further the understanding of the neural systems and the neural representation of stimuli in perception and cognition (Andrillon and Kouider, 2020; Hennevin et al., 2007; Peigneux et al., 2001; Velluti, 1997).

Finally, we acknowledge several limitations of the study. Some of the findings could describe general aspects of auditory processing in different states of consciousness, but it would be naïve to take the results *prima facie* as generalizable. First, the modulation of tone-pairs neural similarities between conscious states were observed in the specific context of the experimental design used here. The auditory modality exhibits remarkable context-dependencies such as behavioural settings, attentional level and task-specific information, which can greatly modulate neural activity in response to sounds (Bekinschtein et al., 2009; Kuchibhotla and Bathellier, 2018; Nelken, 2020). Furthermore, context-dependent auditory processing is observed even in responses to changes in the range of pure tones (Garrido et al., 2013; Regev et al., 2020; Stilp and Assgari, 2019). Therefore, it is likely that the results reported here were shaped by parameters such as the range of tone frequencies, the adaptor tone frequency, and possibly by different adaptation dynamics in wakefulness and sleep. Thus, the observed auditory neural similarity modulations should not be taken as absolute values but as influenced by the specifics of the experimental design. Second, to maximize the number of trials, stimuli were presented at a rate of ∼2Hz, which overlapped with sleep specific features. Slow waves which are in the range of 0.5-4Hz, and K-complexes which lasts 0.5-2 seconds, fall in the same frequency range as stimuli presentation rate. Therefore, even if advantageous for trials number, this high presentation rate prevents a clear separation of the contribution of specific features of each sleep stage to the auditory responses, and makes it difficult to separate the influence of slow waves, spindles and K-complexes on the neural similarity between tones. Third, the EEG resolution provided the required temporal sensitivity to capture the dynamical changes along processing time but limited the ability to infer which brain areas are involved in this functional reorganization. The precise auditory pathway and the underlying mechanism of sensory remapping across the sleep-wake remain to be revealed by future studies.

To conclude, sleep takes centre stage as a model to understand the mechanisms of neural representations of perception and functional reorganization of the brain between conscious states (Andrillon and Kouider, 2020; Mensen et al., 2019). Here, by recordings whole-brain neural activity using high-density EEG during a full night’s sleep, we capture different aspects of sensory processing across the sleep-wake cycle, and provide converging evidence for state-dependent functional reorganization. Precisely how our conscious state shapes auditory processing depends on the relationship between the particular characteristics of the stimulus and neural processes. These findings stress the importance of a systematic investigation of different axes in the sensory map as well as a range of contexts to uncover the rules by which sleep reshapes sensory processing.

## Methods

### Full night sleep EEG experiment

#### Participants

Thirty-eight participants (20 women; age = 25.1 ± 4.96 years mean ± standard deviation [SD]) were recruited to the study and gave written informed consent to procedures approved by the University of Cambridge Research Ethics Committee, in accordance with the Declaration of Helsinki. Participants received monetary compensation for taking part in the experiment. Inclusion criteria were normal hearing, and no history of neurological, psychiatric or sleep disorder. Out of the 38 participants, one participant was excluded due to a technical problem with earphones and another participant due to insufficient sleep time. Data from a total of 36 participants was therefore retained for the analysis.

#### Stimuli

Ten pure tones synthesized in Matlab (2012b) were presented binaurally using Etymotics earphones, at a supra-threshold volume comfortable to the participant. Tone frequencies were 30% apart spanning a range from 500Hz to 5302Hz (500, 650, 845, 1098, 1428, 1856, 2413, 3137, 4079 and 5302 Hz). Two additional tone frequencies (6893Hz and 8961Hz) were presented only to the first five participants and were therefore excluded from the analysis. Each tone was played for 100 ms, with a 10 ms fade-in/fade-out of the sound, and Inter-trial interval (ITI) of ∼500 ms, jittered between 480 and 520 ms, and against a pink noise background (1/f noise) which is known to improve sleep stability (Zhou et al., 2012).

#### Auditory paradigm

Participants listened to a pattern of auditory stimuli including pairs of pure tones. The first tone in every pair was an ‘Adaptor’ (A) tone of 500Hz and was presented in order to “tune” the brain to a common baseline tone, and create a common context for all tones (Sankaran et al., 2018). The second tone in a pair (T) was one of the 10 pure tones mentioned above. Each A-T pair was repeated for 10 times and the ten repetitions created a *mini-block*. 10 mini-blocks presented in a random order created a *block* (Fig. 1). In each mini-block one tone in the sequence was omitted. A wakefulness session was composed of 24 blocks, accumulating in 2400 “A” trials and 216 “T” trials for each tone frequency. During the wakefulness sessions, at the end of every block (∼2.5 min), the experimenter asked the participant to verbally rate a statement from the Amsterdam Resting-State Questionnaire (Diaz et al., 2013). This task was designed to maximize the likelihood that participants would remain awake throughout the session. During the sleep session, auditory stimuli were presented while participants had no task to perform and the number of blocks and trials depended on each individual’s sleep duration (Table 1 and 2).

### Experimental procedure

Participants arrived at the EEG lab at a pre-selected time based on their usual sleep schedule (∼21:00). After the experimental procedure was explained and written informed consent was obtained, participants were seated in a shielded chamber of the EEG room and a 128-channel EEG net was applied on their head (Electrical Geodesics Inc system). The experimental procedure included an auditory paradigm presented during a 1-hour pre-sleep wakefulness session, and full-night sleep session (Table 2). During the wakefulness session, participants sat on a chair in a dim and soundproof room, and were instructed to keep their eyes closed while the auditory stimuli were presented. In the following sleep session, the auditory paradigm was initiated several minutes after the lights were turned off, when participants were comfortably lying in bed, and continued until they woke up in the morning.

### EEG acquisition and pre-processing

The EEG signal was recorded with a 128-channel Sensors using a GES 300 Electrical Geodesic amplifier, at a sampling rate of 1000 Hz (Electrical Geodesics Inc system/Philip Neuro). Conductive gel was applied to each electrode to ensure that the impedance between the scalp and electrodes was kept below 70 kΩ. Peripheral electrodes on neck, cheeks, and forehead were excluded from the analysis due to potential high movement-related noise, retaining 92 electrodes over the scalp surface (Chennu et al., 2014). EEG data was band-pass filtered between 0.5 and 40Hz, segmented in epochs of 550ms (from -100ms pre-stimulus to 450ms post-stimulus), and down-sampled to 250Hz. Next, noisy electrodes were removed if the average signal was above or below 3.5 SD. Eye movements and noise artefacts were removed by means of an Independent Component Analysis (Delorme and Makeig, 2004). Epochs containing voltage fluctuations exceeding ±150 μV in wakefulness or ±300 μV in sleep were also excluded. A liberal threshold was set in sleep to avoid epoch exclusion due to slow waves and K-complexes. Then, the data was re-referenced to the common average of the signal and bad electrodes were interpolated. EEGLAB MATLAB toolbox (version 9.2) and Python (version 3.6) were used for data pre-processing.

### Sleep scoring

Two independent experienced sleep examiners blind to stimuli onset/offset times, scored offline 30 s-long windows of EEG data according to established guidelines (Iber et al., 2007). The two scoring lists were subsequently compared and controversial epochs were inspected again and discussed until an agreement was reached. EEG and EOG signals were first re-referenced to mastoids and then EEG signals were bandpass filtered between 0.1 and 45Hz, EOG between 0.2 and 5Hz. EMG signals were obtained from local derivation and were high-pass filtered above 20 Hz.

### Analysis

#### Auditory evoked potentials analysis

ERPs were computed as the average across five centro-frontal electrodes (E6, E13, E112 E7, E106 in 128-channel EGI net) selected based on (Duncan et al., 2009), for each participant, tone frequency and conscious state. The number of trials differ between sleep stages, in accordance with each participant’s individual sleep architecture (Table 1). Thus, to avoid biases in the results, we equalized the number of trials by randomly sampling for each state and tone as many trials as the ones present in the condition with the smallest number of trials. The procedure was repeated for each participant 1000 times, and then the EEG signal was averaged across the 1000 random samples. Differences in ERPs between conscious states were first investigated by means of a cluster permutation analysis. T-tests conducted between all possible pairs of states (i.e., wakefulness-N2, wakefulness-N3, wakefulness-REM, N2-N3, N2-REM, N3-REM) identified 11 significant clusters (FDR corrected for multiple comparisons). Then, to further characterise the relation between tone properties and conscious state, a linear mixed-effects model analysis was applied to each cluster, where *tone frequency* and a *state* were modelled as fixed effects, while the variable *participant* was treated as a random effect (i.e., random intercept). Given that the assumptions of normality and homoskedasticity of the residuals were violated, the variances were explicitly modelled, by mean of the varIndent function provided of the nlme package in R. This made residuals normal (Anderson-Darling’s A < 1.04, p > 0.05 Bonferroni corrected), and homoscedastic (Levene’s F < 1.175, p > 0.186) for all models. Nonparametric dependent samples effect sizes were calculated as Wsr = Z/sqrt(n) (Rosenthal et al., 1994), where **Z** is the Wilcoxon signed-rank statistic and **n** is the sample size.

#### Similarity magnitude analysis

To estimate the similarity between brain activity in response to different tone frequencies in each of the conscious states, we calculated a similarity measure that we termed ‘*Tone similarity Index*’. Specifically, at each state, for each tone and each electrode, we computed an ERP as the average of all the available trials weighted by their standard deviation (SD). This normalization was done in order to account for the difference in number of trials for different tones, subjects, and states. The neural similarity between each of the 36 tone-pairs was estimated by Spearman’s coefficient between normalized ERPs in each electrode. The obtained correlation coefficients were then averaged across all 92 included electrodes, generating a *Tone Similarity Index* per tone-pair, state and participant. To obtain normally-distributed data and control for multicollinearity, Tone similarity index values were transformed by means of a Fisher z-transformation (i.e., an inverse hyperbolic tangent function), and centred to the grand mean. Next, the Tone Similarity Index across all tone-pairs and participants was computed for a rolling time window of 60 ms (progressively shifted of one time point (=4ms) at each step), which allowed us to obtained a temporally dynamic representation of the Tone Similarity Index values for each state (Fig. 3a). This analysis showed that the Tone similarity index is fluctuating along the auditory processing time. Using cluster permutation analysis (p < 0.01, threshold-free cluster enhancement), we identified two main time windows characterised by differences in Tone Similarity Index between states were observed: an early (12-236 ms) and a late processing time window (256-448 ms). Thus, to avoid masking of one processing stage by another, all the subsequent analyses were conducted for the early and late processing time windows separately.

Next, a linear mixed-effects model analysis was applied to evaluate whether the magnitude of Tone Similarity Index in each participant (36) was influenced by tone-pair (36), conscious state (4), and time window (2) using R nlme package (Version 1.2.5033). We considered several different possible ways of modelling the Tone Similarity Index and compared the evaluated models by means of a Likelihood Ratio test (Table 3) following a protocol outlined in (Zuur et al., 2009). Model fits were estimated by using the Restricted Maximum Likelihood (REML) method when comparing models which differed in their random effects, and the Maximum Likelihood (ML), when comparing models which differed in their fixed effects. The best fitting model was the one with the largest Log likelihood, and it presented conscious *state*, *tone-pairs* and *time window* as fixed effects (all interactions included, with the exception of the triple one), and the variable *participants* as random intercept; also a random slope for *state* was included in the model (Table 3). The variance in the data was opportunely modelled by means of the varIdent function of the R nlme package. To uncover the nature of the observed interactions, additional linear mixed-effects model analyses were performed on subsets of the data, for each time window (Tables 4 and 5), each conscious state, and pairs of state separately. A Tukey’s correction was applied to account for the multiple comparisons and two-tail tests were performed.

#### Representational Similarity Analyses

Representational Similarity Analysis (RSA) (Kriegeskorte et al., 2008), particularly suitable to detecting second-order isomorphisms (Shepard and Chipman, 1970), was applied to test whether different conscious states presented a comparable pattern of Tone Similarity Indexes. First, individual 9 by 9 *Representational Similarity Matrices* (RSMs) were generated generated, where each column and each row corresponded to one of the presented tones, and each cell of the matrix contained the measured neural similarity between the ERPs elicited by each pair of tones, i.e., the Tone Similarity Index value. We obtained distinct RSMs for each participant, at each conscious state and for each temporal window. Two RSA analyses were conducted. First, we ran an *item-analysis*, where we estimated the correlations between RSMs averaged across participants (Fig. 5a-d), and assessed their significance by means of Mantel’s tests. Subsequently, we ran a random-effects analysis (i.e., group-level analysis), where each participants’ RSMs were correlated between pairs of states, within the same temporal window. The hypothesis that the correlations were different from zero was tested by means of bootstrapping (10000 repetitions). A Sidak correction was applied to account for the multiple comparisons.

To characterise the Tone Similarity Indexes structure in each conscious state, we estimated how they related to three conceptual models. The first model is the *Frequency Difference Model,* which is based on the physical distance in Hertz between tone frequencies. In this model, the relations between tone-pairs were calculated by subtracting the lower tone frequency from the higher tone frequency (Fig. 7a). The second model is the *Greenwood Model*, which is derived from studies on the cochlear structure, and relates the anatomic location of the inner ear hair cells to the tone frequencies at which they are stimulated (Greenwood, 1990). Here, the relations between tone-pairs were calculated by means of the Greenwood function from which we obtained the distance in millimetres along the basilar membrane between locations that are maximally excited by each tone frequency. The Greenwood function is:

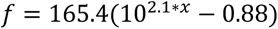

Where *f* is the frequency stimulating the ear, and x is the proportion of total basilar membrane length. The Greenwood function was then opportunely inverted and rearranged so that we could obtain the distance in mm between the two points on the basilar membrane that were excited by each pairs of tones:

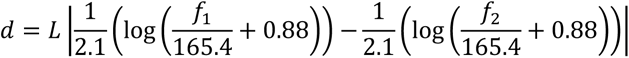

Where *L* is the length of the basilar membrane (=35mm), and *f*_1_ and *f*_2_ are the frequencies of the considered pair of tones.

The Third model is the *Mel* Mode, which we developed based on the Mel scale, and represents the non-linear relationship between perceived pitch and tone frequency (Micheyl et al., 2012; Moore, 2003; Stevens et al., 1937). We computed the relations between tone-pairs by dividing the Mel value corresponding to the higher frequency by the Mel value corresponding to the lower frequency. Specifically, we converted each tone frequency (650, 845, 1098, 1428, 1856, 2413, 3137, 4079, 5302Hz) into its Mel value using the following set of formulas:

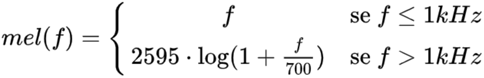

All models described above expressed the relationship between tone-pairs in terms of distance (dissimilarity), rather than similarity. Thus, we first converted participants’ RSMs into Representational Dissimilarity Matrices (RDMs) by replacing each cell value with the *Tone Similarity Index* with the term *1-Tone Similarity Index*. Then, we estimated the Spearman’s correlation between the three model matrices and participants RDMs, for each state and time window separately. The significance of the correlations was estimated using a bootstrapping procedure. The significance *α* level was corrected for multiple comparisons, controlling for the familywise error rate with a Sidák correction.

Next, linear mixed-effects model analysis was applied to evaluate whether the predictive value of the models for each participant (36) was influenced by conscious state (4), and time windows (2) (Table 6).

#### Hierarchical clustering analysis (Dendrograms)

Hierarchical clustering analysis was used to represent the difference in similarity magnitude (Fig. 6a-b) and the difference in similarity pattern (Fig. 6c-d) between conscious states and between tones in each conscious state (Fig. 6e-l). For the magnitude analysis, a 4 by 4 dissimilarity matrix was computed, where each row and column corresponded to one of the four conscious states, and each cell presented the mean effect size (Cohen’d) of the differences in similarity magnitude between pairs of states, normalised by their standard deviation. In the pattern analysis, each cell of the dissimilarity matrix presented the dissimilarity value 1-Spearman’s r, where the Spearman’s r was the Tone Similarity Index as computed from the RSA item-analysis described in the previous section. Hierarchical clustering consists of an iterative method. At each iteration, within a dissimilarity matrix, it identifies the two clusters (i.e., the two matrix cells) that are closest to each other. Once the closest clusters are found, they are merged into a single cluster *i* and the dissimilarity matrix is re-computed, replacing the rows and columns of the two original clusters with one representing the newly formed cluster. The distances between the new cluster *i* and the remaining clusters are therefore estimated and the process is repeated until only one cluster remains. We initially considered each conscious state as an individual cluster and, at each step, identified the closest clusters by means of a Nearest Point Algorithm. A similar procedure was used to obtain the dendrograms representing the relationship between tones in each conscious state (Fig. 6e-l). This time, the input dissimilarity matrices were the average RDMs per state obtained from the RSMs represented in Figure (4a-d) (with RDM = 1-RSM) MATLAB open source software FieldTrip (Oostenveld et al. 2011), and Python (version 3.6) and R (version 1.3.1073) customized scripts were used for ERP and similarity analysis.

## Acknowledgments

This work was supported by the Blavatnik family Foundation (to A.A.), Royal Society—Kohn International fellowship (NF150851 to A.A.), European Molecular Biology Organization (EMBO) fellowship (ALTF 33-2016 to A.A.), and The Wellcome Trust (WT093811MA to T.A.B. and UNS49938 to A.A.). We are grateful to Antje Ihlefeld, Emily Coffey, Israel Nelken, Antje Heinrich, Daniel Bor and Andres Canales-Johnson for their comments and for fruitful discussions.

## Author contributions

Identification, A.A. and T.A.B.; Experimental Design, A.A. and T.A.B.; Experiments: A.A., A.L., A.K., and D.N.; Data Analysis: A.A., C.T. and T.A.B.; Manuscript Writing, A.A., C.T., and T.A.B.

